# Morphological and temporal variation in early embryogenesis contributes to species divergence in Malawi cichlid fishes

**DOI:** 10.1101/2022.09.16.508246

**Authors:** Aleksandra Marconi, Cassandra Yang, Samuel McKay, M. Emília Santos

**Author notes:** Authors for correspondence &.

## Abstract

The cichlid fishes comprise the largest extant vertebrate family and are the quintessential example of rapid “explosive” adaptive radiations and phenotypic diversification. Despite low genetic divergence, East African cichlids harbour a spectacular intra- and interspecific morphological diversity, including the hyper-variable, neural crest (NC)-derived traits such as colouration and craniofacial skeleton. Although the genetic and developmental basis of these phenotypes has been investigated, understanding of when, and specifically how early, in ontogeny species-specific differences emerge, remains limited. Since adult traits often originate during embryonic development, the processes of embryogenesis could serve as a potential source of species-specific variation. Consequently, we compared the features of embryogenesis between three Malawi cichlid species – *Astatotilapia calliptera, Tropheops* sp. ‘mauve’ and *Rhamphochromis* sp. ‘chilingali’ – representing a wide spectrum of variation in pigmentation and craniofacial morphologies. Our results showed fundamental differences in multiple aspects of embryogenesis that could underlie interspecific divergence in adult adaptive traits. Firstly, we identified variation in the somite number and signatures of temporal variation, or heterochrony, in the rates of somite formation. The heterochrony was also evident within and between species throughout ontogeny, up to the juvenile stages. Finally, the identified interspecific differences in the development of pigmentation and craniofacial cartilages, present at the earliest stages of their overt formation, provide compelling evidence that the species-specific trajectories begin divergence during early embryogenesis, potentially during somitogenesis and NC development. Altogether, our results expand our understanding of fundamental cichlid biology and provide new insights into the developmental origins of vertebrate morphological diversity.

**Research highlights:** This work details the early development of three divergent Lake Malawi cichlids. A comparative analysis reveals anatomical and timing differences during embryogenesis and indicates divergence of species’ morphologies prior to their overt formation.

## 1 Introduction

The cichlid fishes are a quintessential example of rapid “explosive” adaptive radiation and phenotypic diversification (Genner and Turner, 2005; Henning and Meyer, 2014; Kocher, 2004; Meyer, 1993; Meyer et al., 1990; Salzburger, 2018). Among cichlid radiations, the most species-rich are the multiple radiations in the Great Lakes of East Africa - Victoria, Malawi, and Tanganyika - where hundreds of species evolved in a remarkably short time span (Kocher, 2004). Despite the relative genomic homogeneity (Kocher, 2004; Loh et al., 2008; Malinsky et al., 2018; Moran and Kornfield, 1993), cichlids harbour a spectacular intra- and interspecific diversity in physiology, morphology, behaviour and ecological specialisation, rendering them an attractive model system in a wide range of research fields.

Recent years have brought major efforts towards elucidating the genetic and developmental basis of cichlid morphological diversity. Akin to other vertebrates, a considerable proportion of cichlid phenotypic variation involves structures derived from a common progenitor cell population - the neural crest (NC) (Bronner and LeDouarin, 2012; Bronner and Simões-Costa, 2016). These embryonic multipotent cells arise at the dorsal side of the forming neural tube and migrate away to often distant regions of the embryo where they differentiate into a plethora of cell types. Among these are the elements of the peripheral nervous system, pigment cells and craniofacial cartilages and bones (Douarin and Kalcheim, 1999). The NC-derived phenotypes that received most attention in cichlids are their distinctive pigmentation patterns (Albertson et al., 2014; Brzozowski et al., 2012; Hendrick et al., 2019; Kratochwil et al., 2018, 2022; Liang et al., 2020; Roberts et al., 2017; Santos et al., 2014), and the dramatic variation in craniofacial morphologies, associated with the divergent trophic strategies (Albertson and Kocher, 2006; Conith et al., 2018; Powder and Albertson, 2016; Powder et al., 2014, 2015).

Insights into the genetic basis of cichlid traits have been possible due to their experimental tractability, including viability of hybrid crosses and amenability to genetic manipulations such as CRISPR-Cas9 (Albertson and Kocher, 2006; Clark et al., 2022; Juntti et al., 2013; Kocher, 2004; Li et al., 2021; Powder and Albertson, 2016). Despite these advances, the cellular and developmental mechanisms underlying cichlid morphological diversification remain unknown. Amid the unanswered questions lies the one of when, and specifically how early, in ontogeny do species-specific differences emerge?

Adult morphologies often originate during early embryonic development, hence the processes of embryogenesis could serve as a potential source of species-specific variation. Since NC development (and thus origins of both pigmentation patterns and craniofacial skeleton) coincides with the processes of gastrulation, neurulation and somitogenesis in teleosts (Rocha et al., 2020), how conserved are these stages among cichlids? What are the species-specific and species-generic (i.e. shared by the clade) features of embryonic development?

Species-specific differences could result from variation in embryonic morphology at these early stages of ontogeny, as well as from differences “to the timing or rate of developmental events” (i.e. heterochronies) (Alberch et al., 1979; McKinney and McNamara, 1992). Temporal variation could be especially consequential if involving the period when the processes of embryogenesis (e.g. neurulation and somitogenesis) directly influence NC development (e.g. NC cell migration) and, more indirectly, the subsequent formation of its derivatives.

Here, we compare the embryonic development of Malawi cichlids to assess for potential sources of phenotypic variation. To date, the general features of embryonic development have been described in a handful of African cichlids (Balon, 1977; Fujimura and Okada, 2007; Hendrick et al., 2019; Jones, 1972; Jong et al., 2009; Woltering et al., 2018), yet only a few studies have explicitly compared developmental variation in early ontogeny between or within species (Jones, 1972; Jong et al., 2009), with most being limited to a specific morphological trait (e.g. Hendrick et al., 2019; Powder et al., 2015).

Consequently, we characterise and compare the features of embryogenesis between three Malawi cichlid species – *Astatotilapia calliptera* (AC), *Tropheops* sp. ‘mauve’ (TM) and *Rhamphochromis* sp. ‘chilingali’ (RC) – representing a wide spectrum of morphological variation in pigmentation and craniofacial shape (Figure 1a). TM has a characteristic ‘flattened’ head shape of an algae grazer, RC has the elongated narrow jaws of a pelagic predator and AC shows an intermediate phenotype of an omnivore generalist. The differences in body colouration involve distinct pigment pattern motifs, from vertical bars in TM, horizontal stripes of RC to melanic patches in AC, with the latter comprising features of both bars and stripes. To examine the developmental variation at both intra- and interspecific levels, we have included two populations of *A. calliptera* diverging in body colouration (main lake ‘Salima’ and riverine ‘Mbaka’; Figure 1a-b) in our study system. The ‘Mbaka’ fish of both sexes are noticeably darker than their conspecifics from the ‘Salima’ population. Craniofacial skeleton and pigmentation aside, the characterisation and comparison of developmental processes between these species could be of interest for future morphological evolution studies due to their positions in the Malawi cichlid phylogeny. Notably, AC is thought to strongly resemble the prototype species of the entire radiation in terms of its ecology and phenotype (Malinsky et al., 2018).

**Figure 1.**
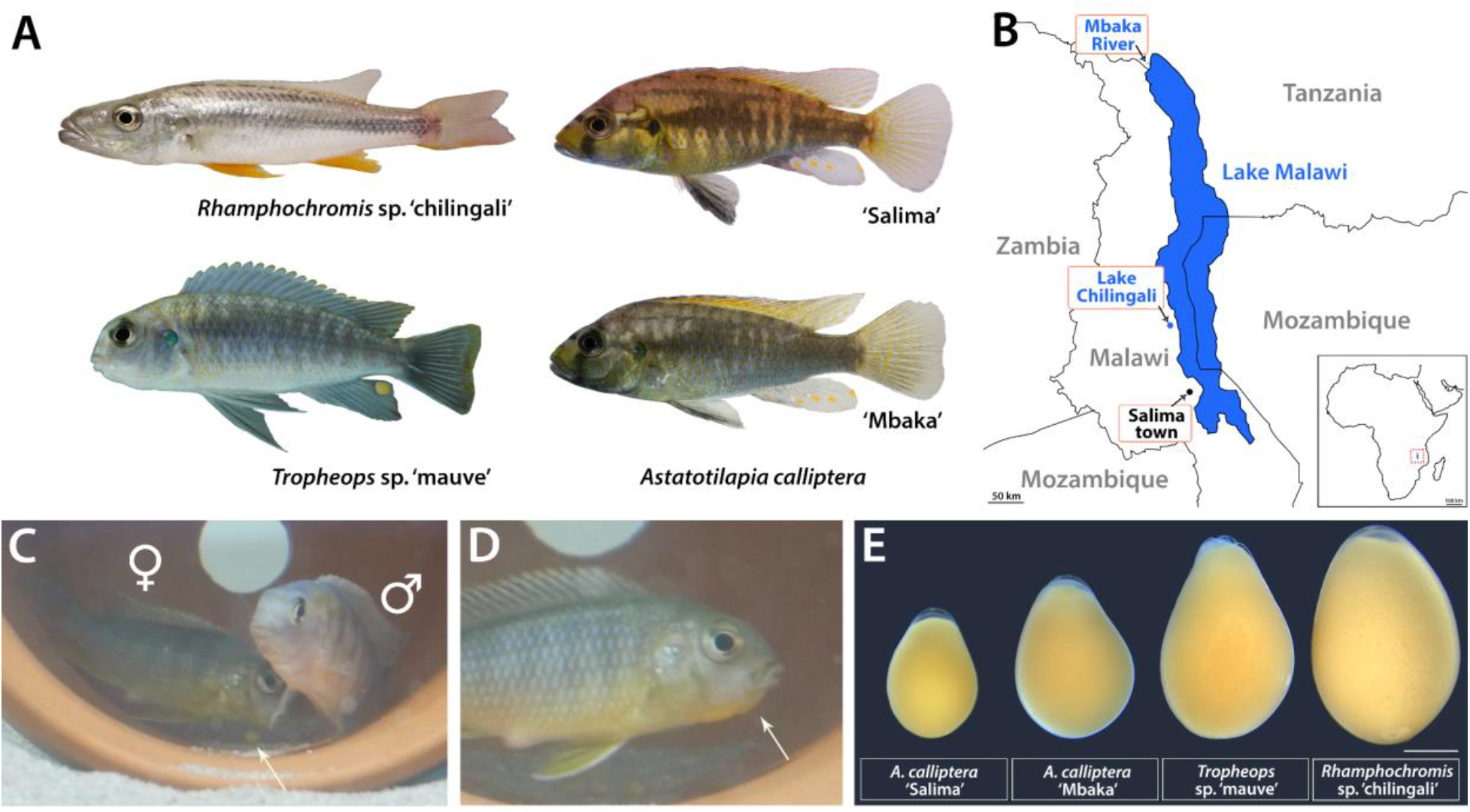
Lake Malawi cichlids. (A) The four focal species of the study are characterised by clear variation in body colouration and craniofacial morphologies. *Rhamphochromis* are pelagic predators of other fish and arthropods and *Tropheops* are algae grazers, whereas the generalist *A. calliptera* is considered to closely correspond to the common ancestor of the Malawi radiation (Malinsky et al., 2018). Note difference in hue between the two morphs of *A. calliptera:* ‘Salima’ inhabiting the main lake reservoir and riverine ‘Mbaka’. All individuals depicted are adult males; (B) All cichlids in the study are endemic to the Lake Malawi basin in East Africa, including Lake Chilingali and River Mbaka, located in close proximity to the main lake. Geographical boundaries drawn after Google Earth 2022; (C) *Tropheops* male and female pair during their courtship behaviour. Note an egg underneath the female (white arrow); (D) Mouthbrooding female with a characteristic protruding ‘chin’ (white arrow). (E) Diversity of egg sizes across the study species, implicating potential variation in the maternal provisioning. Scale bar in E = 1 mm.

Here, we first present an overview of the development of these four cichlid fishes as a staging system. We also examine in more detail the timelines associated with early embryogenesis occurring concomitantly with the NC development (e.g. somitogenesis). Finally, we summarise the earliest stages of the formation of the craniofacial skeleton and compare the timing and order of appearance of three pigment cell types contributing to the adult colouration (black melanophores, reflective iridophores and yellow-orange xanthophores) altogether addressing the question of how early these NC-derived traits diverge in overt morphology. In addition to advancing our understanding of the timing of major developmental events and morphological divergence in early ontogeny, the resulting staging series will provide a valuable addition to the growing interest in cichlid evolutionary developmental biology and facilitate effective experimental design, including comparisons with other teleost systems.

## 2 Materials and methods

### 2.1 Animal husbandry and embryo culture

*Astatotilapia calliptera, Rhamphochromis* sp. ‘chilingali’ and *Tropheops* sp. ‘mauve’ were kept under standardised conditions (26±1°C on a 12:12 hour light cycle). These species are mouthbrooders. To minimise the influence of maternal care, eggs were removed after fertilisation. Since mating took from 30 min to 1.5 hours, we considered the time of fertilisation as approximately within the 1-hour window from the first laid egg. Eggs were cultured individually in 1 mg/L of methylene blue (Sigma Aldrich) in water in 6-well plates (ThermoFisher Scientific) placed on an orbital shaker moving at slow speed at 27°C. All experiments were conducted in compliance with the UK Home Office regulations.

### 2.2 Staging system

Our staging (Supplementary Table S1) is based on the tables for *Astatotilapia burtoni* (Woltering et al., 2018), *Oreochromis niloticus* (Fujimura and Okada, 2007) and *Danio rerio* (Kimmel et al., 1995). We followed the definition proposed by Kratochwil et al. (2015) to measure epiboly as “the ratio between distances between the animal pole and blastoderm margin, and between the animal and vegetal pole”. Embryo age is given in days, counting from the day of fertilisation (day 0).

### 2.3 Imaging of live animals and fixed embryos

Adult photographs (Figure 1) were acquired with a Sony α6600 with a Sony E 30mm f/3.5 lens. For each species and stage reported in the staging (Figures 2–5) and head pigmentation development (Figure 10) series, several embryos (n ≥ 5) from different clutches were examined and followed daily. For stages following hatching, live animals were imaged following anaesthetisation with MS-222 (800 mg/L, Sigma).

**Figure 2.**
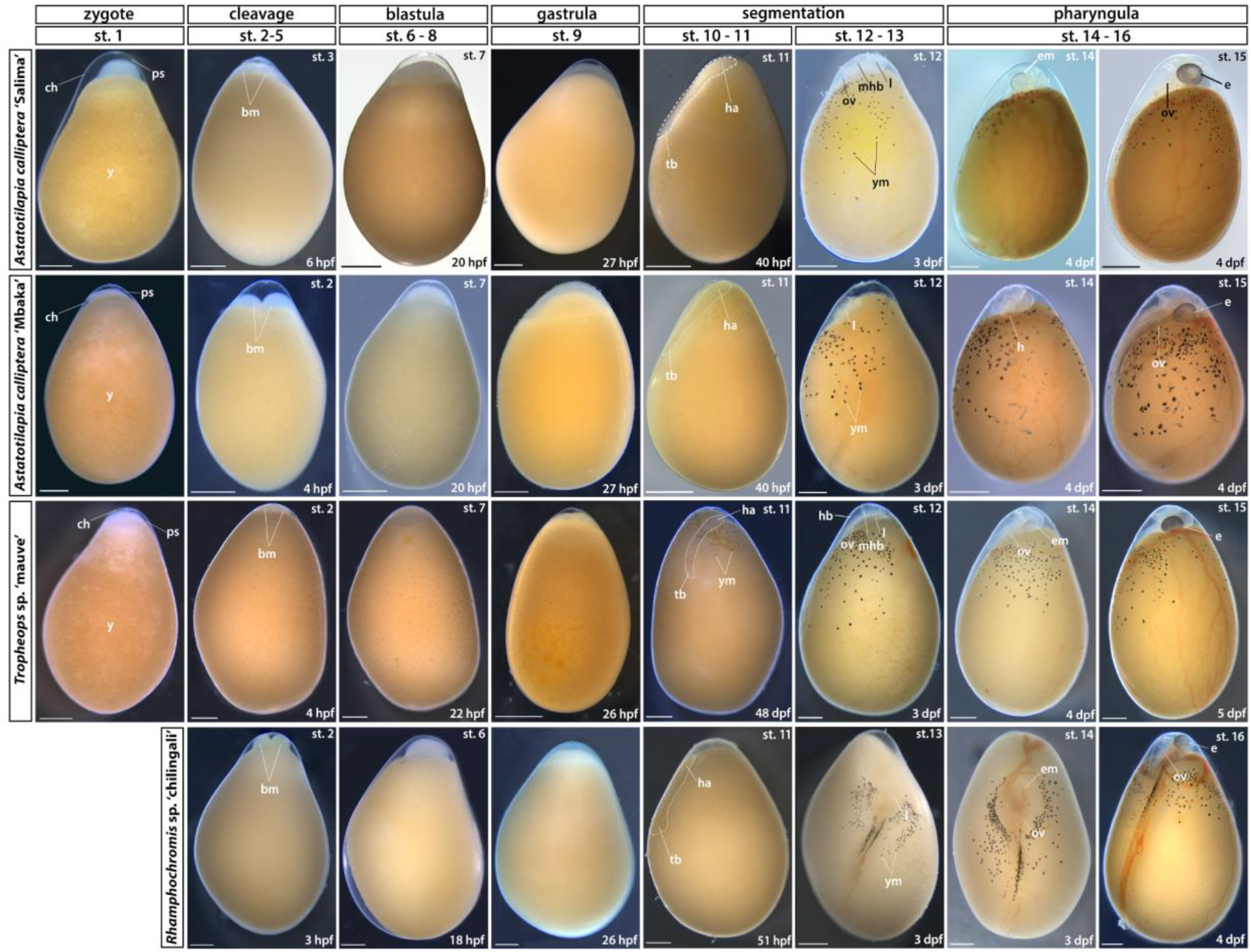
Early embryonic development (zygote to pharyngula). Stage numbering following the staging table of *O. niloticus* (Fujimura and Okada, 2007) (see Supplementary Table S1 for stage descriptions and associated developmental landmarks). Embryos undergoing somitogenesis are outlined in the ‘segmentation’ stage. Lateral views except for dorsal views in RC st. 12-16. No st. 1 (zygote) image available for RC due to their prolonged courting and breeding behaviour. bm – blastomeres; ch – chorion; dpf – days post-fertilisation; e – eye; em – eye melanophores; ha – head anlagen; hb – hindbrain; l – lens; mhb – midbrain-hindbrain boundary; ov – otic vesicle; ps – perivitelline space; st – stage; tb – tailbud; y - yolk; ym – yolk melanophores. Scale bar = 1 mm.

**Figure 3.**
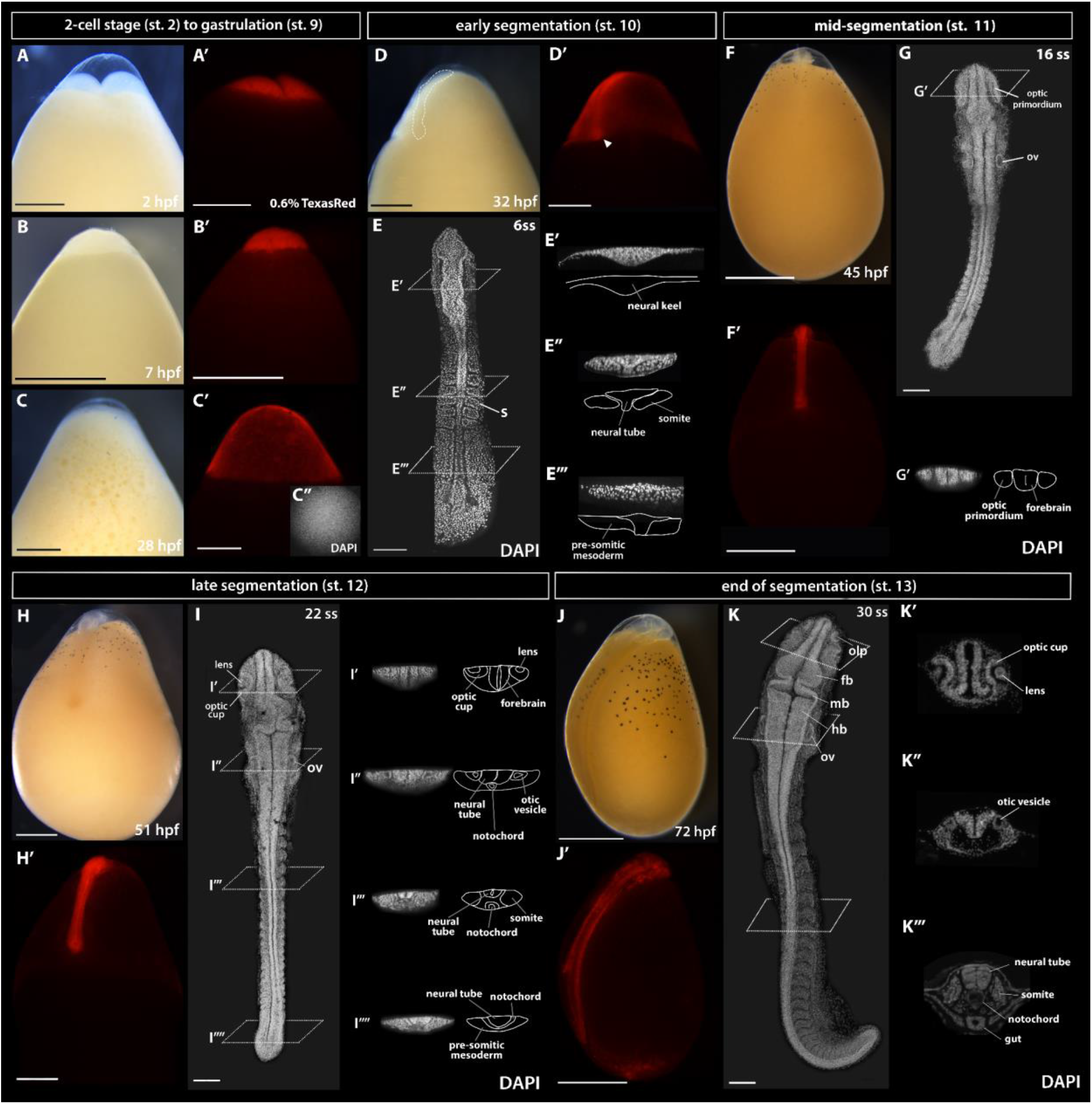
The development of *Astatotilapia calliptera* embryo from 2-cell stage until pharyngula. To observe morphology of live embryos, the fluorescent TexasRed dye was injected at the single-cell stage and it remained detectable in the tissues until at least 3 dpf (pharyngula stage). All whole-mount images (panels A-D’; F-F’; H-H’ and J-J’) are lateral views with the animal pole facing up and the vegetal pole facing down. The same embryo is shown across the series in both brightfield and under RFP fluorescence conditions except for the 2-cell stage (panels A-A’), for which an uninjected control embryo is shown in brightfield (panel A). The embryos in panels E, G, I and K were dissected from yolk, stained with DAPI to visualise nuclei and imaged from the dorsal side with the anterior end of the embryo facing up. The specimen in C’’ was stained with DAPI following removal of the chorion, here shown in the view from the animal pole. Optical sections in all except for 28ss where histological sections are presented. The timing of development is given in hours post-fertilisation (hpf) at 27±1°C. fb - forebrain, hb - hindbrain, l - lens, mb - midbrain, oc - optic cup, olp - olfactory placode, op - optic primordium, ov - otic vesicle, s - somite. Scale bar in A-D’; F-F’; H-H’ and J-J’ = 1 mm; 100 μm in all others.

**Figure 4.**
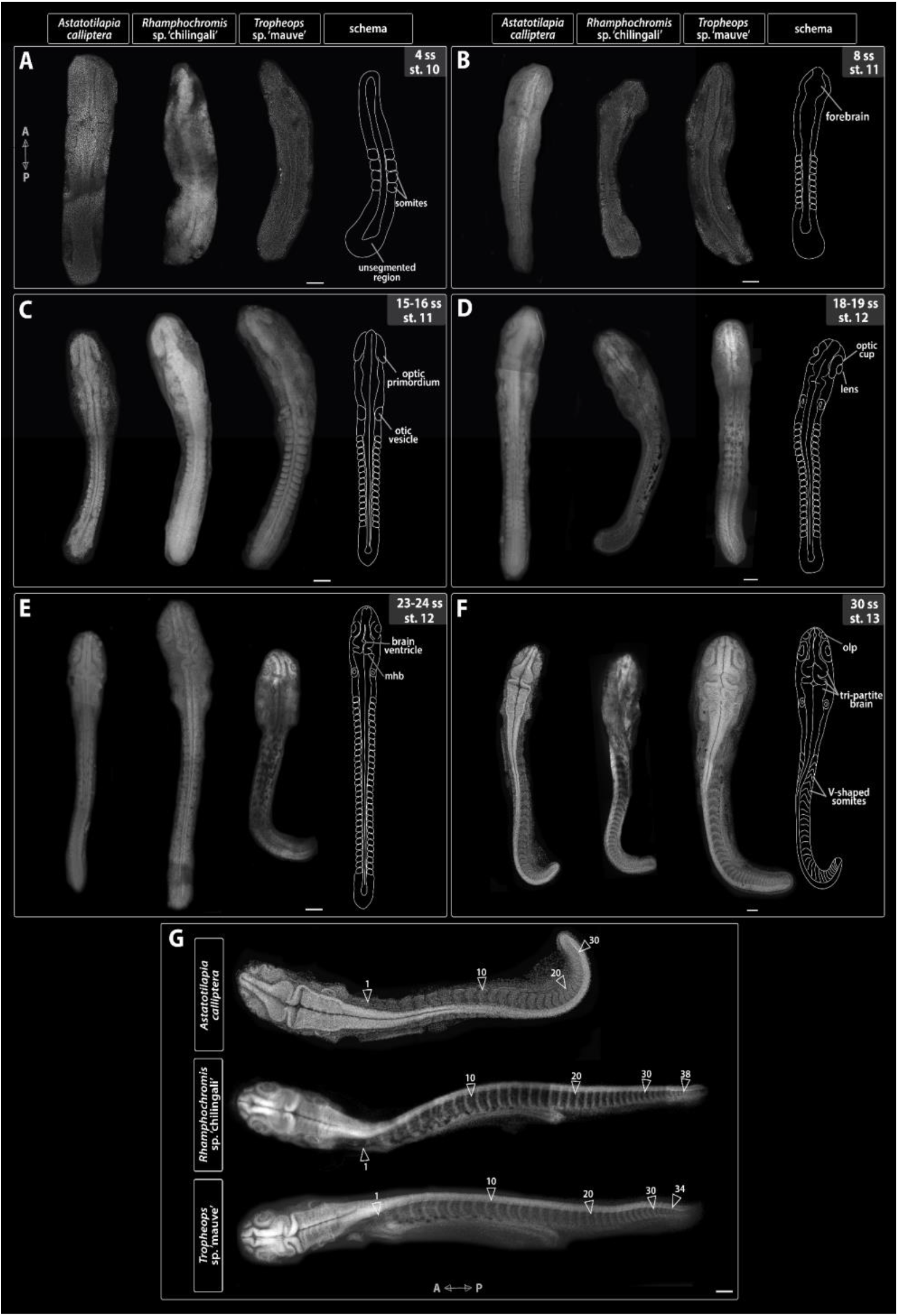
Embryo morphology throughout the segmentation period (st. 10-13). (A-F) Development of anatomical landmarks is correlated with progression of somitogenesis in all examined species i.e. somite stage is a good predictor of embryo morphology. (G) Embryos dissected at the end of their segmentation exhibit differences in the total somite number. All embryos were dissected from yolk, stained with DAPI to visualise nuclei and imaged from the dorsal side with anterior-posterior orientation as indicated on panel A for A-F and in G. A - anterior; mhb - midbrain-hindbrain boundary; olp - olfactory placode; P - posterior; st - stage. Scale bar = 1 mm.

**Figure 5.**
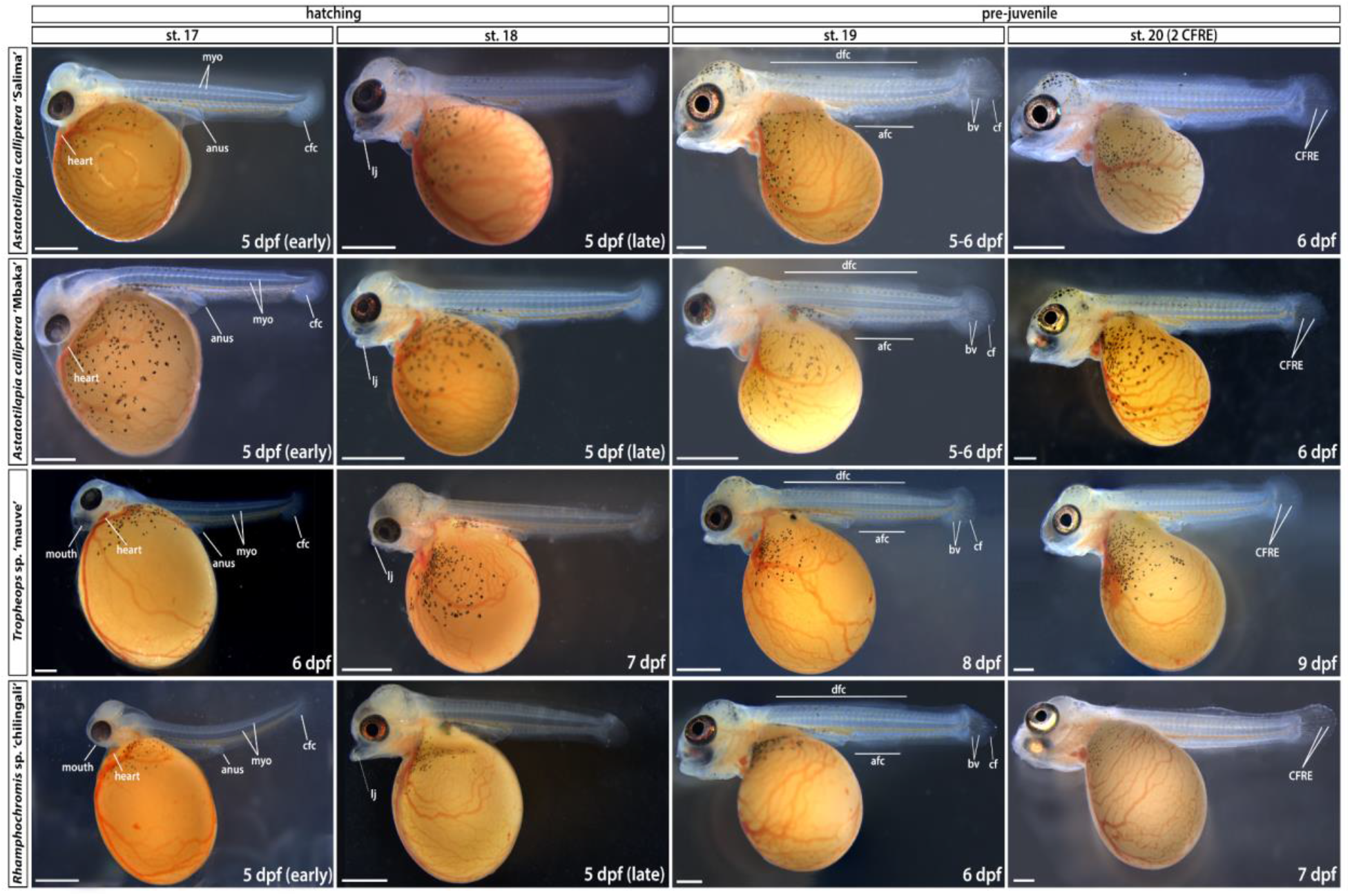
Late embryonic development to early pre-juvenile stages (hatching to stage 20). Stage numbering following the staging table of *O. niloticus* (Fujimura and Okada, 2007) (see Supplementary Table S1 for stage descriptions). afc – anal fin condensation; bv – blood vessels; cf – caudal fin; cfc – caudal fin condensation; CFRE – caudal fin ray elements; dfc – dorsal fin condensation; dpf – days post-fertilisation; myo – myomeres; st – stage. Scale bar = 1 mm.

**Figure 6.**
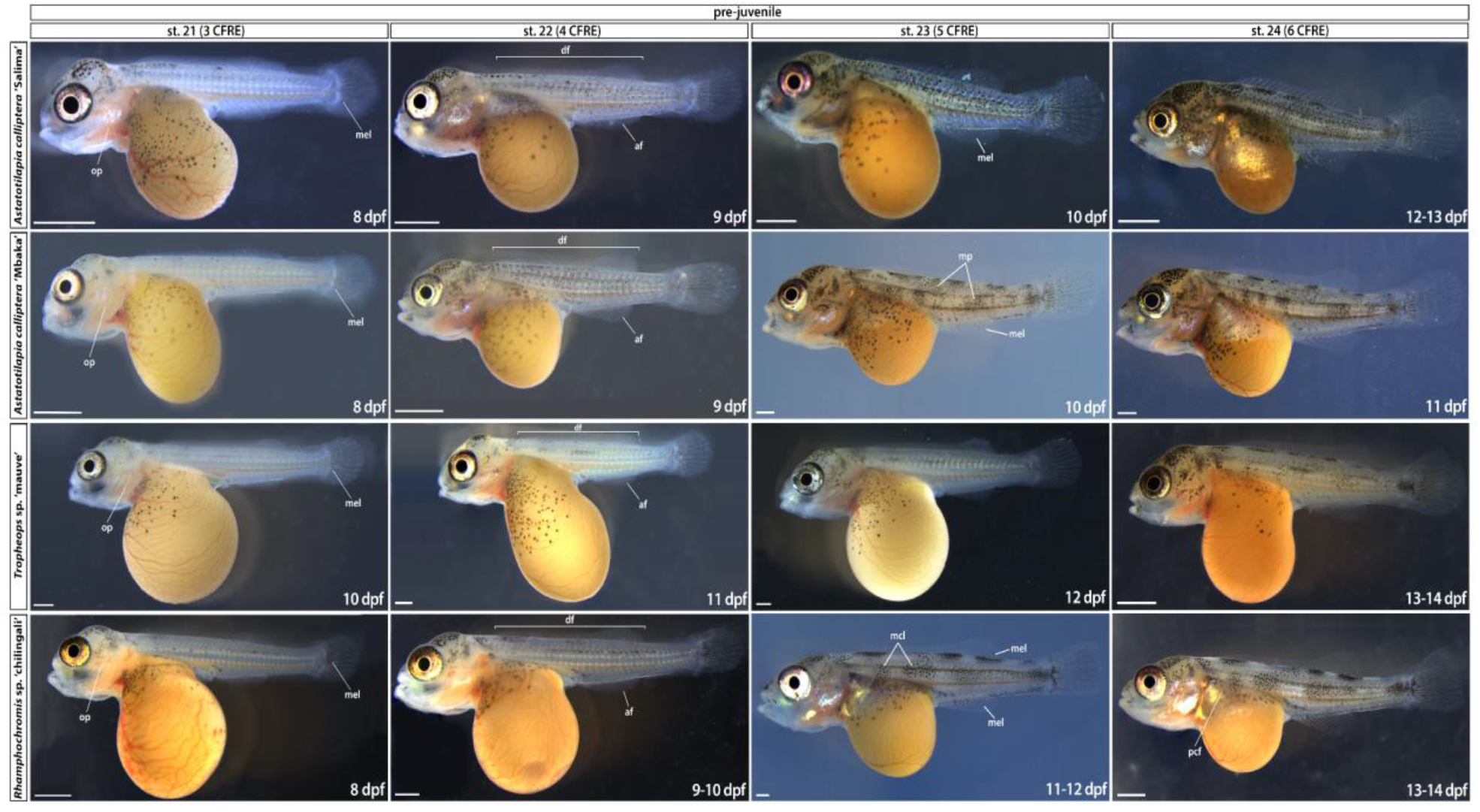
Pre-juvenile development (stages 21 to 24). Stages are delimited based on the number of CFRE. Time ranges (in days post-fertilisation) given to indicate the duration of the corresponding stage. af – anal fin; df – dorsal fin; dpf – days post-fertilisation; mel - melanophore; mcl - melanophore clusters; mp - melanic patches; op – operculum; pcf – pectoral fin; pvf – pelvic fin; st - stage. Scale bar = 1 mm.

**Figure 7.**
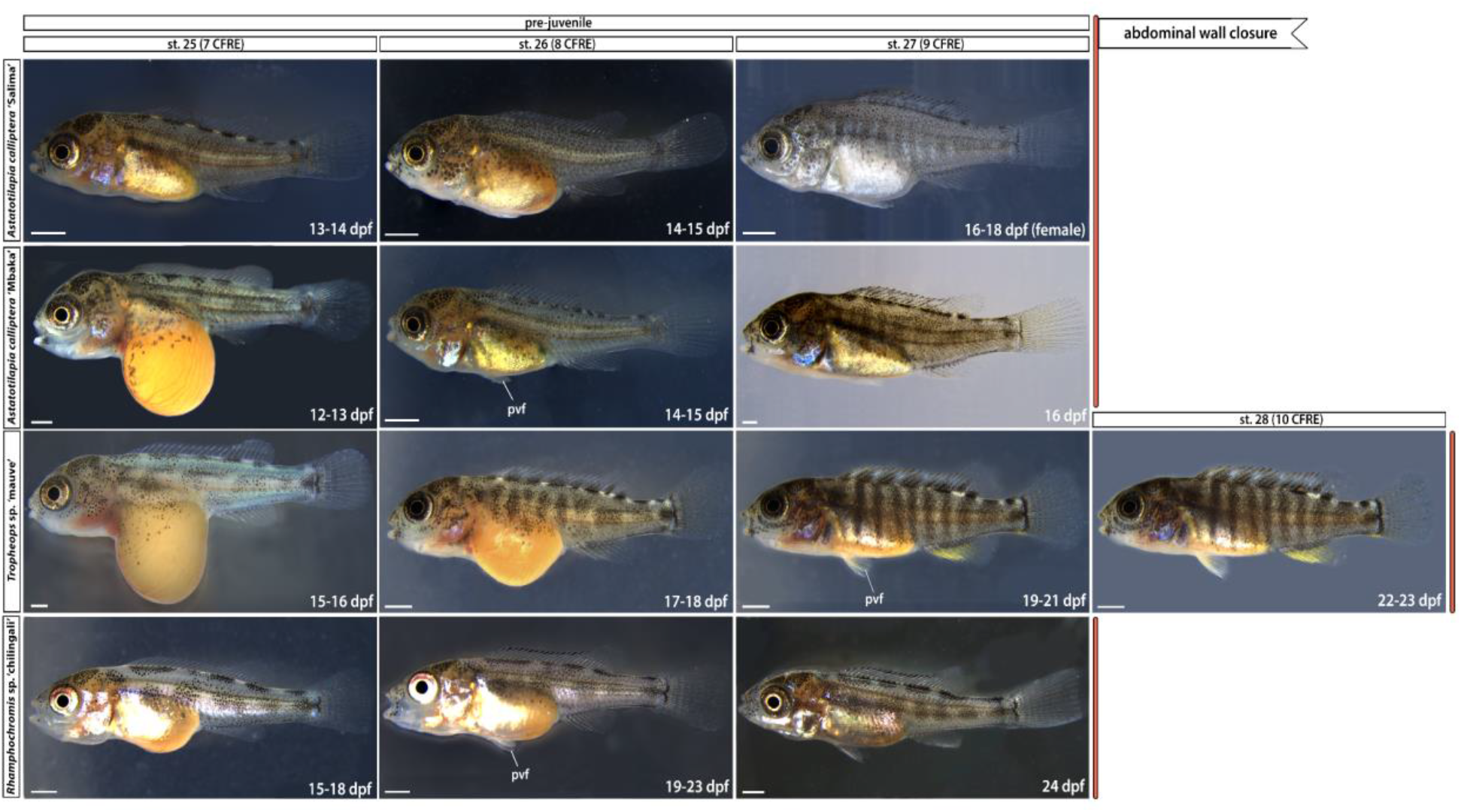
Late pre-juvenile development (stage 25 to abdominal wall closure). Time ranges (in days post-fertilisation) given to indicate the duration of the corresponding stage. Note the silvery hue of *Astatotilapia calliptera* ‘Ssalima’ female at st. 27 compared to a darker male fish shown for st. 26. dpf – days post-fertilisation; pvf – pelvic fin; st – stage. Scale bar = 1 mm.

**Figure 8.**
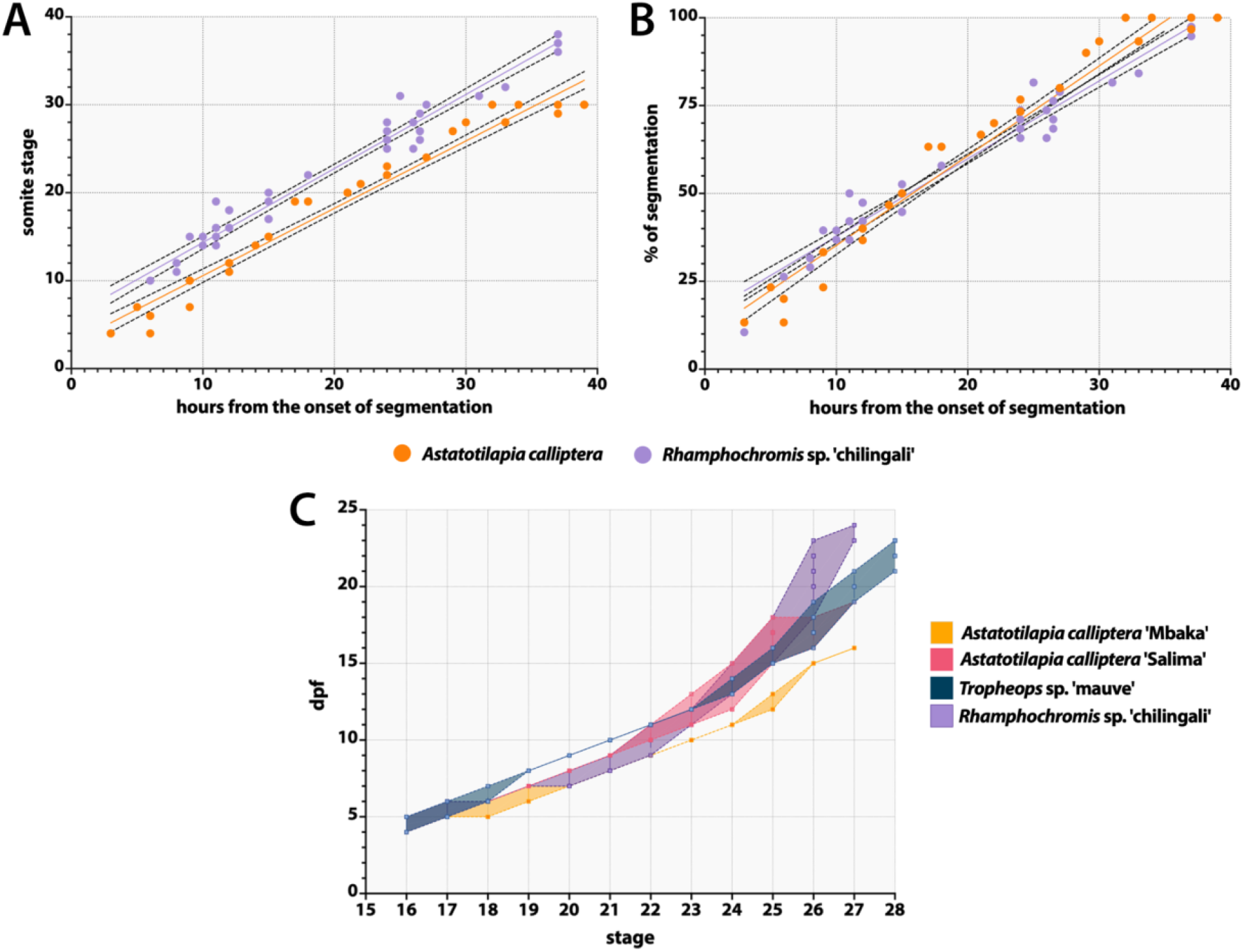
Timelines of cichlid development during the segmentation (st. 10-13) and developmental trajectories from hatching (st. 16) until complete abdominal wall closure (st. 27/28). (A) Somite stage (ss) against hours elapsed from the onset of segmentation. (B) percentage value of the relative completion of segmentation against the hours from the onset of segmentation. Solid trend lines represent an idealised rate of somitogenesis and dotted bands correspond to 95% confidence intervals. (C) Time ranges of the post-hatching stages (st. 16-28). The shaded bands encompass all individual trajectories of the embryos followed through development at 27°C. At least 3 animals from 3 different clutches were inspected at each time-point. Although all raw data points are included, in some instances they are occluded due to the overlap between them.

**Figure 9.**
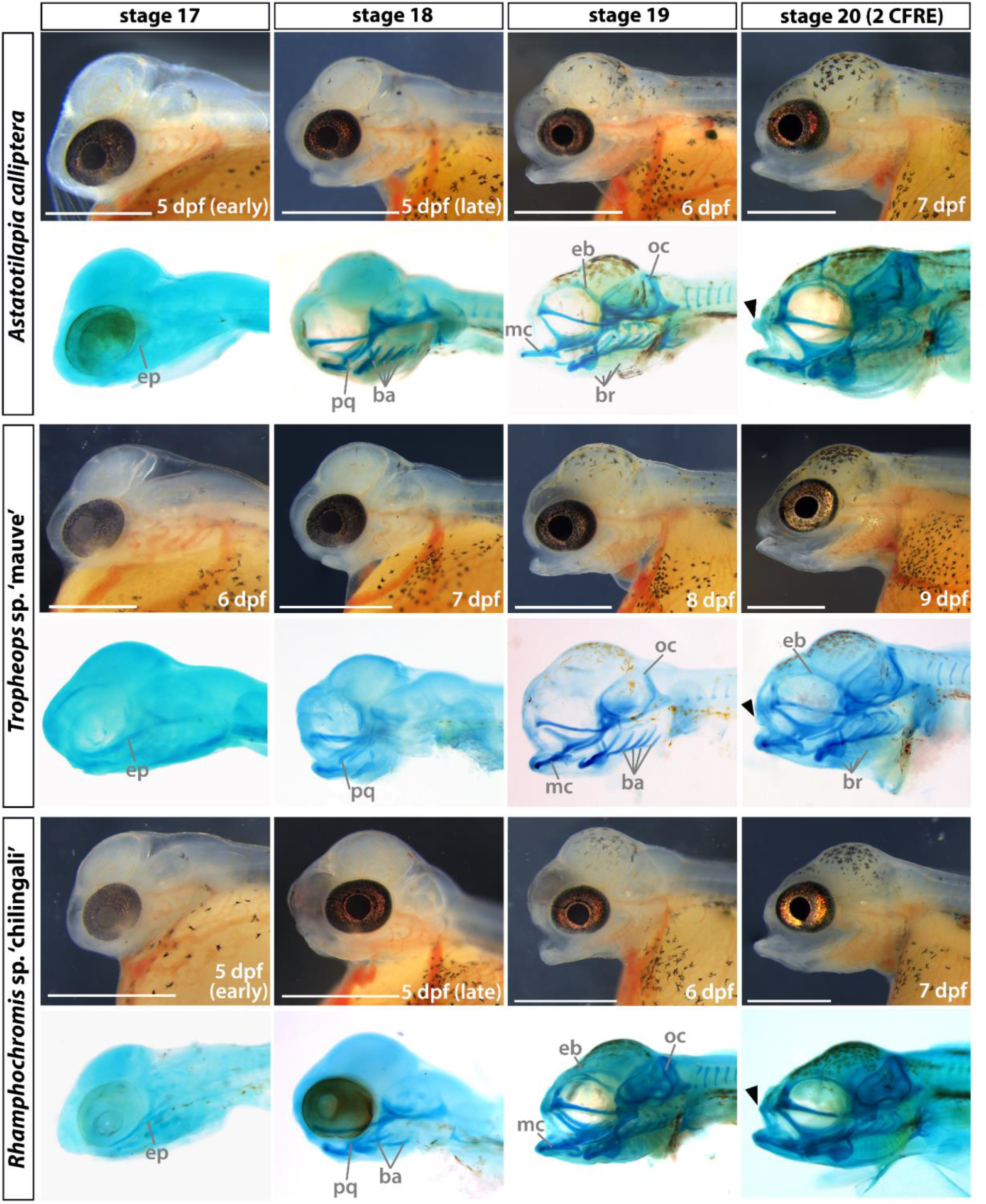
Development of the craniofacial morphology of *Astatotilapia calliptera, Tropheops* sp. ‘mauve’ and *Rhamphochromis* sp. ‘chilingali’ embryos. All images are left lateral views, dorsal side towards the top, rostral side to the left. For each species in turn, panels in the bottom rows present cartilage stains of stage-matched specimens to those depicted in brightfield (top rows). Due to the lack of perceivable differences between AC ‘Salima’ and ‘Mbaka’, only the latter is shown. ba - branchial arches; br - branchiostegal rays; CFRE - caudal fin ray elements; eb - epiphyseal bar; ep - ethmoid plate; mc - Meckel’s cartilage; oc - occipital arch; pq - palatoquadrate. Scale bar = 1 mm.

**Figure 10.**
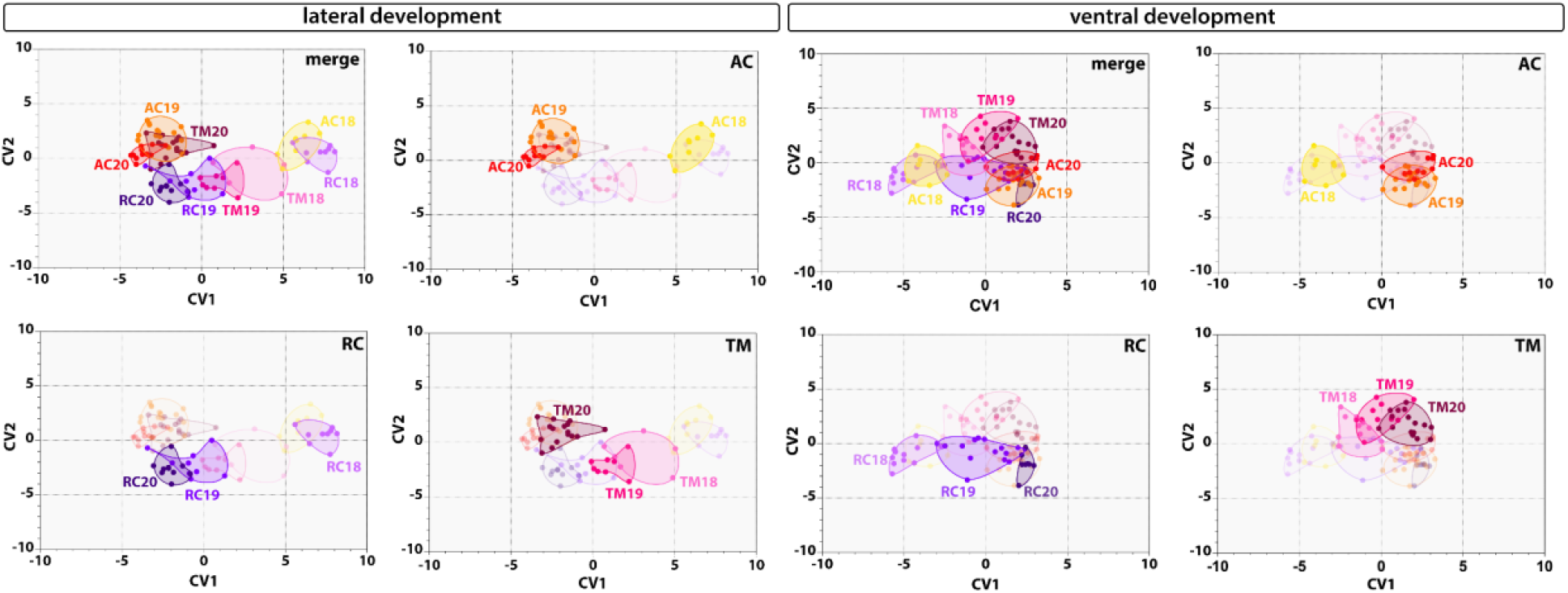
Developmental trajectories of lateral and ventral craniofacial aspects are species-specific. Canonical variate analysis (CVA) through ontogeny shows significant differences in mean shape between species at most stages of development of craniofacial cartilages. Ellipses encompass all data points for each group comprising all specimens of a given species at a specific stage. Panels B-D show the same scatter plot as panel A but with different species highlighted in turn. Similarly, panels F-H are the same as panel E. AC - *Astatotilapia calliptera*, RC - *Rhamphochromis* sp. ‘chilingali’ and TM - *Tropheops* sp. ‘mauve’. Stages are indicated by the number following the species acronym.

To observe the morphology of early embryos more closely, 0.6% v/v solution of dextran labelled with TexasRed (3,000 MW, ThermoFisher Scientific) was injected at one-cell stage in AC eggs using a microinjector system (Applied Scientific Instrumentation). All live embryos (including those injected with TexasRed) were placed in glass-bottomed dishes (Cellvis) in 0.5% low melting point agarose (Promega) for imaging in brightfield or under RFP fluorescence (Figure 7). Images were taken through water to eliminate glare using a Leica M165FC and a Leica DFC7000T camera. Cameras were colour balanced with a grey card (Grey White Balance Colour Card 24 by gwbcolourcard.uk). Using Adobe Photoshop 2022, multiple focal planes of stereoscopic images were aligned and merged, and any background imperfections were removed.

Due to the limited access to one-cell TM and RC embryos, an alternative approach was applied: for each time point, embryos were dechorionated and dissected from yolk and fixed overnight at 4°C in 4% paraformaldehyde (PFA) in 1X phosphate buffered saline (PBS). Samples were rinsed in 1X PBST (PBS+0.01% Tween-20) (20 mins/rinse, twice) and stained with 10nM DAPI (ThermoFisher Scientific) in 1X PBST overnight at 4°C. The embryos were rinsed twice with 1X PBST and mounted on glass-bottomed dishes (Cellvis) with ProLong™ Gold Antifade Mountant (ThermoFisher Scientific). A similar approach was applied to AC embryos used to determine the rates of somitogenesis. DAPI stainings were imaged with an Olympus FV3000. Confocal micrographs were stitched using the Olympus FV3000 software and processed with Fiji (Schindelin et al., 2012) to produce optical sections, collapse z-stacks and adjust image brightness and contrast where necessary. All images were processed for background imperfections in Adobe Photoshop 2022.

### 2.4 Histological sections

Embryos were cleared with histosol (National Diagnostics) (20 min/wash, 3 times) at room temperature and transitioned into wax in a 1:1 molten paraffin:histosol solution (30 min/wash, twice) and placed in molten paraffin (RA Lamb Wax – Fisher Scientific) at 60°C overnight. Molten paraffin was then changed five times (each change lasting >1h) before the tissue was transferred into a Peel-A-Way embedding mould (Sigma) for transverse sectioning. The embedded blocks were left to cool overnight and sectioned using a Leica RM2125 RTS microtome. Sections were mounted on Superfrost plus slides (VWR). The paraffin-embedded sections were dewaxed in histosol (5 min/rinse, twice), 100% EtOH (5 min/rinse, twice) and grading into water through a series of descending EtOH concentrations (90%, 70% and 50%, 5 min/rinse), followed by a final rinse in water (5 min). The slides were coverslipped with Fluoromount G containing DAPI (Southern Biotech) and cured overnight before imaging. The fluorescent micrographs were taken with Zeiss Axioscope A1 and combined into figure plates in Adobe Photoshop 2022.

### 2.5 Cartilage preparations

Specimens were fixed overnight at 4°C in 4% PFA in 1X PBS and dehydrated in increasing increments of EtOH:PBS (20%, 50% and 70%; 10 mins/wash) and stored in 70% EtOH:PBS at −20°C. The embryos were then transferred directly into 30% glacial acetic acid in ethanol and incubated for 2h. Next, they were washed in Alcian Blue solution (0.02% Alcian Blue in acetic ethanol) for 2h and incubated overnight in acetic ethanol. Next, the embryos were stepwise rehydrated from 30% acetic ethanol via 70%, 50%, 25% EtOH:diH_2_O (15 mins/wash). The samples were then bleached in 2% KOH:3% H_2_O_2_ solution to remove skin pigmentation until melanophores turned from black to brown (2-16h) and placed into 0.01% Alizarin Red in 1% KOH for 2h. The specimens were cleared in 3:1 1% KOH:glycerol solution for 1-3 days, depending on the size of the animal. The solution was changed daily until the samples were sufficiently clear. Samples were subsequently transferred to 1:1 solution of 1% KOH:glycerol for 24h and then placed in 1:3 solution of 1% KOH:glycerol until all Alizarin red cleared from non-ossified tissues (replaced with fresh solution daily, 1-3 days). Finally, the specimens were transferred to 80% glycerol for imaging and storage at 4°C. All washes were done with rocking and at room temperature. All cartilage preparations were carried out on at least six separate stage-matched individuals. The specimens were positioned in 100% glycerol in glass-bottomed dishes (Cellvis) and imaged using a Leica M165FC with Leica DFC7000T camera. The images were colour-corrected in Adobe Photoshop 2022.

### 2.6 Geometric morphometrics

To analyse the lateral and ventral development, the positions of homologous anatomical landmarks were collected from images using TPSUtil and TPSDig2 (Rohlf, 2010) following modified landmark protocol of Powder et al. (2015) (Supplementary Figure S1). MorphoJ (Klingenberg, 2011) was used to perform a generalised Procrustes analysis on the landmark coordinate data to exclude any other sources of variation than shape. This software was also used to generate covariance matrices and perform a principal components analysis (PCA). All specimens were staged following definitions detailed in Supplementary Table S1 to avoid the potentially confounding effects of developmental heterochrony when using solely chronological age. We analysed the samples as follows: 1) among stage-matched individuals across all species (i.e. at single developmental time point) and 2) along the ontogeny for each species individually (st. 18-20). To identify differences among these groups, we used canonical variate analysis (CVA) across 10,000 iterations per comparison using Mahalanobis distances.

## 3 Results

### 3.1 Overview of the early development of Malawi cichlids

To provide a visual guide assisting embryo staging for further analyses, we present an overview of development from fertilisation until early juvenile stages, focusing on the formation of the major features of the external morphology. The consistent staging nomenclature (Supplementary Table S1) used throughout this study makes it the first comparative analysis of the entire embryogenesis across multiple closely related, yet morphologically distinct cichlid species.

Overall, the development of the examined species closely resembles descriptions for other African cichlids (Fujimura and Okada, 2007; Jong et al., 2009; Morrison et al., 2001; Otten, 1981; Saemi-Komsari et al., 2018, Woltering et al., 2018). As such, and unlike zebrafish or medaka, these cichlids have large and yolk-rich eggs, supplying essential nutrients for the developing embryo until it transforms into an actively feeding juvenile (Figure 1e). The eggs are surrounded by a translucent chorion and the embryo develops on top an opaque yolk (ch and y in Figure 2, st. 1).

#### 3.1.1 Embryonic development from zygote to gastrula

On the first day post-fertilisation (dpf), the embryos undergo meroblastic cleavage divisions and enter the blastula period. The first mitotic division occurs within 2.5 hours post fertilisation (hpf) and each following one is paced at 2-3h (Figure 2, st. 2-5). The blastomere (bm in Figure 2, st. 2-5) arrangement resembles that of zebrafish, with regular grids of 2×2, 2×4, 4×4 and 4×8 forming on the animal pole of the egg. By the 64-cell stage, the individual blastomeres are difficult to distinguish and the regularity of cell arrangement is no longer discernible. Over the next few hours, the cells start to form a ball-shaped blastodisc located on the animal pole of the egg which subsequently flattens as embryo progresses from blastula (Figure 2, st. 6-8) to gastrula (Figure 2, st. 9). The incremental flattening of blastodisc transforms it into a uniformly thick layer - the blastoderm - which begins to cover the yolk from the animal pole in the process of epiboly (Figure 2, st. 9-11).

#### 3.1.2 Development from gastrulation through somitogenesis

From the onset of gastrulation (Figure 2, st. 9) until mid-somitogenesis (Figure 2, segmentation, st. 10-12), the embryo proper becomes difficult to distinguish from the surrounding extraembryonic tissues and its features are not easily observable in live embryos (Figure 2, st. 9-12). Consequently, we examined these stages more closely, using fluorescent dye microinjections of single-cell embryos with TexasRed and nuclear DAPI staining of dissected and fixed embryos (Figures 3–4). The period of somitogenesis (i.e. process of sequential addition of mesodermal somites) was of particular interest due to its temporal concurrence with specification and migration of the NC cells (NCCs). We determined the chronology of gastrulation and somitogenesis and the total number of generated somites in each species, since somite stage (ss) is a commonly used index to stage-match embryos. Due to its established experimental amenability (Clark et al., 2022), we used *Astatotilapia calliptera* (AC) embryos to illustrate the common features of cichlid development as observed in live specimens (Figure 3).

Following cleavage and blastula stages (Figure 3a and b, respectively), all species reach 20% epiboly around 26-28 hpf (Figure 2, st. 9, Figure 3c). By 28-30 hpf (25-30% epiboly), the cell density in the blastoderm is no longer uniform, with a more densely occupied region at one side of the blastoderm, marking the future embryonic axis (Figure 3d-d’). The embryo undergoes gastrulation, a process which results in three germ layers. Similarly to other cichlids (Jong et al., 2009; Kratochwil et al., 2015; Woltering et al., 2018) but unlike zebrafish (Kimmel et al., 1995), embryos start segmenting before epiboly is complete (Figure 2, st. 10-13; Figure 3d-e).

By 32 hpf in AC (Figure 2d, st. 10, 4-6 ss; Figure 3e), the embryonic axis is discernible by eye and the posterior end of the embryo (Figure 3e’”) is flatter and wider (plate-like) compared to the anterior end. At this point, although the first few somite pairs have already formed (Figure 3e and e”), the embryo still thins down to a single cell layer (epidermis) at the extreme margins where it joins the rest of the blastoderm spreading over the yolk, as visible in the optical transverse sections at the prospective head region (Figure 3e’). At this early stage in somitogenesis, the region located anterior to the first somites has the characteristic triangular shape of the neural keel (Figure 3e’), a structure formed from neural plate and a precursor of the neural tube in teleosts (Lowery and Sive, 2004).

Over the next 24h, the embryos thicken and elongate via sequential addition of somites. The unsegmented tail region has a bud-like appearance (Figure 3f-i) and progressively shrinks as the somitogenesis progresses. Concomitant with the body axis elongation of, optic, otic and olfactory vesicles (precursors of the eye, ear and the olfactory epithelium, respectively) develop (Figure 2, st. 12-13; Figure 3f-k, Figure 4d-f). Specifically, around 8-9ss (Figure 4b), the optic primordia begin to form from the anterior neural keel, whereas the otic vesicles located beside the caudal region of the hindbrain become discernible by 12ss (Figure 3g; Figure 4c). The first visible pigmentation - black melanophores - appears on the yolk along the mid-section of the elongating embryo and soon after spread over the yolk (Figure 2, st. 10-11 for *Tropheops sp*. ‘mauve’ (TM), st. 12-13 for AC and *Rhamphochromis* sp. ‘chilingali’ (RC)). The lenses in the optic cups are clearly visible by 18ss in all species (Figure 3i; Figure 4d). The brain grows and undergoes regionalisation during the second half of the segmentation period (Figure 2, st. 12-13; Figure 3f-k; Figure 4c-f). The midbrain-hindbrain boundary (the isthmus) of the developing brain becomes prominent around 22-23ss (Figure 2, st. 12-13; Figure 4e). The olfactory placodes and three brain vesicles (forebrain, midbrain and hindbrain) become apparent by 30ss (Figure 3k; Figure 4f). Concurrently with the late phase of somitogenesis (>28ss), epiboly approaches 90% (i.e. the posterior end of the embryo reaches the vegetal pole of the egg) and trunk somites become V-shaped (Figure 3j-k and Figure 4f). At this point, the beat of the transparent heart is visible in embryos dissected from yolk (data not shown).

Although the overt processes of embryogenesis occurring until the end of the segmentation period seem to be generally conserved between the examined species, we observed divergence in the total number of generated somites. Specifically, AC has 30-32 somites, TM 34, whereas RC up to 38 (Figure 4g). Despite this key difference, the developmental progression in these species, marked by the gradual acquisition of anatomical landmarks such as the optic and otic vesicles, seems to be more tightly correlated to the number of already formed somites (i.e. somite stage) than to the relative completion of somitogenesis (the ratio of existing somites to the total number per species). Moreover, except for the addition of new somites and increase in size, we did not observe any further changes to external morphology of both RC and TM past 30ss (i.e. the end of segmentation in AC) when compared to AC (Figure 4g). Altogether, our results show the embryo morphology at a given somite stage is largely matched between species throughout the segmentation period, irrespective of the variation in the overall number of produced somites.

#### 3.1.3 Development during the pharyngula period

The pharyngula period (st. 14-16) is characterised by the progressive development of dark eye colouration, an increasing vasculature on the yolk surface and circulation of red blood cells. The pigmentation of retinal epithelium starts at st. 14 (Figure 2) and increases in intensity until the hatching period (Figure 4, st. 17-18) when the eyes become fully opaque. The somites located posterior to the trunk region gradually change from a rounded rectangular shape to chevron-like Vs (Figure 4g) as they differentiate into myotomes (myo in Figure 5, st. 17) in an anterior-to-posterior order. The embryo now extends around the entire length of the yolk with the tail curling inside the chorion (Figure 2 st. 16, Figure 3j). The head thickens and becomes bulbous with the development of the brain and elements of the face (Figure 2 st. 16, Figure 3j).

#### 3.1.4 Development from hatching (st. 17) to st. 20: early stages of development of skin pigmentation and skeletal system

Hatching period encompasses the transition from pharyngula (a late embryonic stage) to post-embryonic (or pre-juvenile) stages of development and marks the onset of a gradual formation of adult traits including the head cartilaginous skeleton and body pigmentation (Figure 5, st. 17-18). As with other direct-developing species (Balon, 1977, 1999; Jones et al. 1972; Woltering et al., 2018) the adult body plan, including the anal and dorsal fins, is progressively attained throughout the post-embryonic stages (Figures 5–7). Stages 17-19 (Figure 5) were delimited based on the head morphology (Supplementary Table S1), whereas from st. 20 onwards, the number of caudal fin ray elements (CFRE) was used as a diagnostic feature. At st. 17, the ventral side of the head is attached to the yolk but a small opening, marking future mouth, can be distinguished just above the heart (Figure 5). The tail is separated from the yolk sac. Despite some intra- and inter-clutch variability, most embryos hatch at this stage (5 dpf for AC and RC, 6 dpf for TM). At st. 18 (Figure 5), the head lifts up from the yolk, the mouth opens and occasional movements (‘wiggles’) of the tail are observed. At the following stage (Figure 5, st. 19), both the operculum covering the gills and the lower jaw begin to move sporadically and the mesenchymal condensations marking future anal and dorsal fin develop (afc and dfc in Figure 5, respectively). The developing blood vessels in the caudal fin (bv and cf in Figure 5) become more prominent. Embryos at st. 20 (Figure 5) have fully functional and rapidly moving oral jaws and two CFRE. This stage marks the transition from embryonic to pre-juvenile stages when the adult body plan and external morphology will be gradually acquired.

#### 3.1.3 Development from st. 21 to complete body wall closure (st. 27-28): continued formation of species-specific phenotypes

The embryos start to right themselves and soon can swim upright. From st. 21, the caudal, dorsal and anal fins develop pigmentation, starting with melanophores (mel in Figure 6, st. 21). Differences in the head and jaw morphology, body shape and pigmentation patterns between species are increasingly noticeable. For instance, at st. 23 (Figure 6), the melanophore flank pigmentation remains scarce in TM, AC ‘Mbaka’ has irregular melanic patches (mep, Figure 6 st. 23) spread out over its flanks, whereas in RC, melanophores form a specific pattern of large, oval clusters distributed on the flanks along the midline and dorsum. Notably, the melanophore distribution is considerably more uniform across the flank in AC ‘Salima’ compared to its co-specific ‘Mbaka’.

At the same stage, the inter-specific differences in body shape and head morphology are also apparent (Figure 6, st. 23). For example, the body of RC is more elongated along the anterior-posterior axis compared to shorter and more corpulent bodies of TM and AC. The pelvic fins form by st. 26 in all species except TM, in which they appear at st. 27 (pvf in Figure 7). Interestingly, the closure of the abdominal wall over the yolk sac, marking the onset of the juvenile period and thus the end of our staging table, is similarly delayed by one stage in TM compared to AC and RC (Figure 7, st. 27-28). These two instances exemplify heterochrony in the development of morphological features where particular characters do not always appear simultaneously or in the same order between species. At this point, the yolk is fully absorbed into the body cavity and juvenile fish start to feed actively.

Overall, the overview of the early development of Malawi cichlids presented here is a clear illustration of the wide range of biological diversity harboured both between and within species. Based on the readily visible external features alone, the embryonic development of examined cichlids is very similar until hatching (Figure 2 and 4), followed by a rapid appearance of species-specific morphologies (Figures 5–7). Despite these broad similarities in the early ontogeny, the examined species exhibited considerable variation in the timing or rate of development, including during the segmentation period and post-hatching stages.

### 3.2 Pervasive heterochrony during cichlid embryogenesis

#### 3.2.1 Duration of somitogenesis is similar between species despite variation in rates (st. 10-13)

The temporal periodicity of somite addition, termed the segmentation clock, is known to exhibit vast species-specific variation in the pace of its progression (Hubaud and Pourquié, 2014). Considering the differences in the total numbers of formed somites between species (30-32 in AC, 34 in TM and 38 in RC, Figure 4g), we tested for variation in segmentation rates. To collect a representative sample, we aimed to sample 2-3 embryos from the same clutch (n=3) at each given time point. However, due to TM’s small clutch size, we were unable to follow its development in a similarly detailed manner (Supplementary Table S2).

Intriguingly, the first five somites in both AC and RC seem to form simultaneously, potentially explaining the elevated addition rates during early segmentation, which then decrease approaching the end of somitogenesis (Figure 8a). Overall, the rates in both species follow linear trends, with no pronounced intraspecific variation. The slight differences among RC specimens could be explained by the duration of their mating behaviour (2-3h). Although somites form at a faster pace in RC throughout the process, when the inferred rate is scaled proportionally to the species-specific total somite number, the segmentation is completed within a similar time frame in both AC and RC (Figure 8b). Similarly, even though the appearance of morphological structures is tightly coupled with somite addition in both species up to 30ss (i.e. the end of segmentation in AC, section 3.1.2), since segmentation is progressing faster in RC than AC, these anatomies also develop faster in RC and only additional somites are added in the remaining time during somitogenesis. These heterochronic shifts explain how the overall duration of segmentation is conserved between species with different somite numbers.

#### 3.2.2 Post-hatching development (st. 16-28) exhibits intra- and interspecific temporal variability

Since interspecific temporal differences were observed also throughout the post-hatching stages (Figures 5–7), we quantified this variation by following embryos across their development and contrasting their developmental trajectories against chronological time elapsed since the day of fertilisation (Figure 8c). Our results indicate that the development of each of the species follows a slightly distinct temporal path with overlaps at specific stages. For example, the developmental trajectory of AC ‘Mbaka’ diverges around st. 18 from the other species and remains separate except for a brief overlap with RC at st. 20-22. The former develops the fastest among the species, progressing between consecutive stages within a day to reach the last embryonic stage (st. 27) at 16 dpf. Contrarily, RC embryos tend to be the slowest (st. 27 at 23-24 dpf). Although there is one more stage (st. 28) in TM’s trajectory, they still enter the juvenile period ahead of RC (21-23 dpf Figures 7 and 8c). Interestingly, the timing and duration of some stages demonstrate intra- and interspecific variation, particularly in the later phase of the pre-juvenile development (represented by the width of the shaded regions in Figure 8c, st. 22-28). Among the examined species, the embryos of AC ‘Mbaka’ exhibit the least temporal variability, whereas RC embryos vary the most. These findings add further evidence that the developmental trajectories of these cichlids, especially in the temporal aspect, seem to be already distinct at the time of embryogenesis. The considerable variation in timings between species highlights the potential risks associated with relying solely on embryo age or morphological landmarks to guide comparative studies in this clade.

### 3.3 The early ontogenies of skeletal system and body pigmentation

To determine when differences in cichlid NC-derived trait development are first evident, we investigated the early formation of the craniofacial skeleton and skin pigmentation. The overt development of these traits begins around the time of hatching (Figure 5, st. 17-18) and continues throughout the post-embryonic period (Figures 6–7).

#### 3.3.1 I Divergence in craniofacial shape is evident at the onset of cartilage deposition

We investigated the formation of craniofacial cartilages (Figure 9) to assess for qualitative and quantitative differences between species across early ontogeny (Supplementary Figure S1, Supplementary Table S2). The first cartilaginous element of the pharyngeal skeleton - primordial ethmoid plate - begins to form at st. 17 (5 dpf in all species, ep in Figure 9). At st. 18 (5-6 dpf in AC and RC, 7 dpf in TM), in addition to the formation of almost all the cartilaginous structures of the lower jaw (except for the basihyal), the palatoquadrate of the upper jaw is also present (pq in Figure 9). The branchial arch elements, although formed in AC and RC, are not detected at this stage in TM (ba in Figure 9, st. 18). By st. 19 (6-7 dpf in AC and RC, 8 dpf in TM), chondrogenic condensations of the occipital arch appear around the eye orbit and the vomerine process, the epiphyseal bar and the branchiostegal rays have formed. The articular process of the Meckel’s cartilage and basihyal form in the jaw and the branchial arches are now fully developed in TM (ba in Figure 9, st. 19). At st. 20 (7-8 dpf in AC, 7 dpf in RC and 9 dpf in TM), the upper lip (black arrowheads, Figure 9) is present. The consistent temporal shift in the development of the craniofacial complex by at least one day in TM compared to AC and RC (Figure 9) further suggests that timing differences likely contribute to interspecific divergence.

To quantitatively compare divergence in craniofacial development between species, we conducted geometric morphometric analyses on the lateral and ventral views of specimens stained for cartilage taken for st. 18-20 (Figures 10–11, Supplementary Figure S1, Supplementary Table S3). We used canonical variate analysis (CVA) to assess how well sample groups (here defined by species, stages and a combination of the two) can be differentiated from one another by maximising the between-group to within-group variance ratio. The results showed that within each species-specific trajectory, the lateral shapes at different stages occupy largely or entirely distinct morphospaces (Figure 10a-d), with the primary axis of variation (CV1) corresponding to developmental age. The extent of between-stage overlap within each species’ ontogeny was similar among all three taxa. Notably, the largest distance in morphospace between consecutive stages was observed in AC and RC, suggesting a large change to craniofacial shape early in ontogeny in these species compared to a more gradual development in TM. Finally, the trajectories between species were generally distinct, except for the large overlap in morphospaces between AC and TM at stages 19 and 20 respectively.

**Figure 11.**
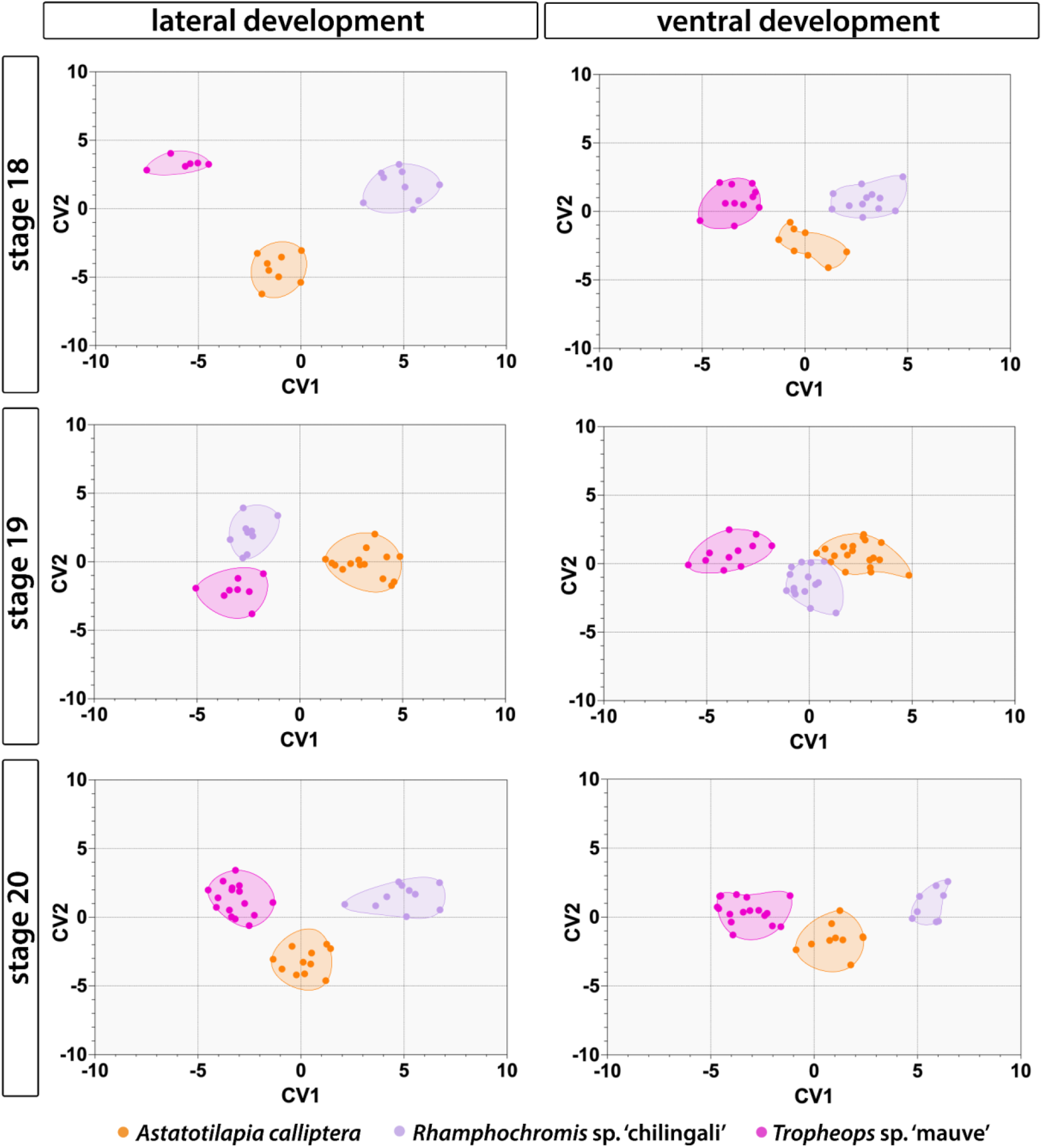
Stage-wise comparison of lateral and ventral morphologies reveals patterns of convergence and divergence between species along the ontogeny. Each panel presents a CVA conducted for all specimens at single developmental time points. Ellipses encompass all data points for each group comprising all specimens of a given species at the indicated stage.

Contrarily to lateral aspect, the overlap between ontogenetic trajectories was considerable across the ventral development, including between AC and RC at st. 18 (Figure 10). However, akin to lateral aspect, the craniofacial shapes of TM were mostly distinct from stage-matched AC and RC (Figure 11), suggesting an early divergence of the developmental trajectory of this species. Intriguingly, the large shape change between stages 18 and 19 in lateral development for AC and RC was not present for the latter in the ventral trajectory (Figure 11). Altogether, our results demonstrate that despite similarities, the morphology of craniofacial cartilages is species-specific from the onset of its overt formation and continues to diverge through time.

#### 3.3.2 I Differences in head epidermal pigmentation are apparent as soon as the first pigment cells appear

To identify species-specific divergence in pigmentation development, we examined the timing of pigment cell appearance on the dorsal side of the head. The head epidermis is the first area to be populated by all three major cell types, appearing significantly earlier than the chromatophores underlying the flank pigment patterns. A summary schematic of the different cell populations and specific head regions described below is presented in Figure 12c.

**Figure 12.**
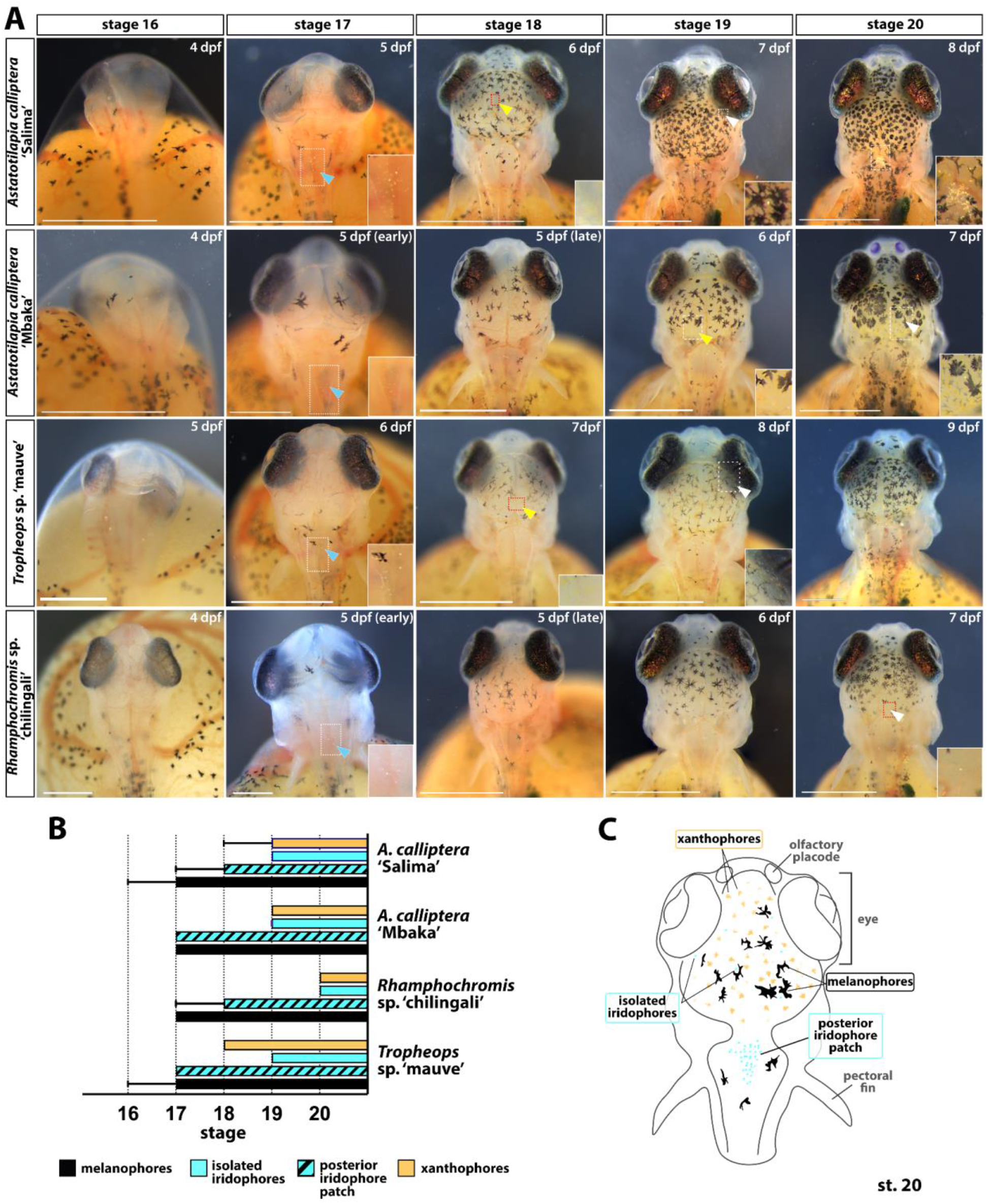
Development of skin pigmentation on the dorsal head region in *Astatotilapia calliptera* ‘Mbaka’ and ‘Salima’*, Rhamphochromis* sp. “chilingali” *and Tropheops* sp. “mauve” embryos. (A) Developmental series of head pigmentation from appearance of the first pigmented cells (st. 16, hatching) to the presence of all three pigment cell types in all studied species (st. 20). Insets show close-ups of the regions bounded by white rectangles. (B) Timeline of pigment cell appearance. Error bars indicate the earliest appearance of a given chromatophore type. Bar indicates presence in all examined specimens. The distinction of two iridophore sub-populations (i.e. ‘unique’ and ‘posterior patch’) as highlighted in C. (C) Schematic summary of the common features of the chromatophore distribution among cichlids in this study as observed at st. 20, viewed from the dorsal perspective. st - stage. At least 3 animals from two different clutches were examined. Scale bar in A = 1 mm.

The first black melanophores are detected at st. 16 (Figure 12a) in some embryos of AC ‘Salima’ and TM but not in the other two species (Figure 12b). At st. 17 (Figure 12a), melanophores on the dorsal surface of the head and rostral trunk region are now present in all inspected individuals. In AC and TM, these are thin and elongated with several projections each, whereas the larger and wider ones are not fully opaque. In contrast, melanophores observed in RC appear small and rounded, presumably due to centralised localisation of the dark pigment in the cell. Round reflective iridophores begin to aggregate in the posterior patch (blue arrowheads, Figure 12a; Figure 12b) in all species but not across all examined embryos. By st. 18 (Figure 12a), more pigment cells have appeared on the dorsal surface of the head in all species, including the first signs of yellow pigmentation in AC ‘Salima’ and TM (yellow arrowheads, Figure 12a, st. 18; Figure 12b). Iridophores continue to accumulate in the skin covering the dorsal hindbrain. Isolated round iridophores are sparsely distributed among the melanophores on the head region in all species except for RC (white arrowheads, Figure 12a, st. 19; Figure 12b). Lastly, at st. 20, all three chromatophore types are present in all taxa (Figure 12a-c) and the melanophores on the dorsal head region tend to have a ‘snowflake-like’ morphology i.e. with a small, circular centre and multiple radially extending projections that make contact between neighbouring cells. Intriguingly, some melanophores (as seen on AC ‘Mbaka’ in Figure 12a, st. 20) do not fit this description and instead of long projections, have jagged edges and appear to cover a larger surface area than the former type. By now, single iridophores (white arrowheads, Figure 12a and c, st. 20) and xanthophores in the posterior region are also found in RC but the latter appear less conspicuous (or fainter) and numerous than in the other taxa. Another major difference in pigmentation at this stage concerns the abundance of iridophores in the posterior patch i.e. it is much more pronounced in AC ‘Salima’ than in any other species (Figure 12a, st. 20). Generally, although the order of appearance of each chromatophore type (melanophores first, xanthophores last) and the stereotypical cell distributions at stage 20 (Figure 12c) showed a close resemblance between species, we observed intra- and interspecific variation in the morphology, localisation, and abundance of the specific cell types relative to the chronological (dpf) and developmental age (stage).

Altogether, our results demonstrate that the initial gross similarities in external morphology present during pharyngula and early hatching periods (st. 14-17, Figures 2 and 5) progressively decrease with the gradual appearance of species-specific phenotypes, including the pigmentation patterns and craniofacial morphologies, as the animals progress into juvenile and adult stages. Considering that both of these phenotypes showed interspecific variation at the onset of their overt formation, we hypothesise that these differences begin to be specified beforehand i.e. during early embryonic development.

## 4 Discussion

A long-standing goal of evolutionary developmental biology is to understand the mechanisms underlying morphological diversification. Since a considerable proportion of the crucial morphogenetic processes occur during embryonic development, studying variation in the early ontogeny is key to understanding when and how divergent phenotypes form. The challenge in addressing these questions lies partly in the paucity of multispecies systems showing natural variation yet offering the experimental tractability for comparative embryology similar to the conventional models. Here, we took advantage of the Malawi cichlid system to compare early development across multiple closely related species harbouring extensive divergence in adult phenotypes such as body colouration and craniofacial skeleton. These adaptively relevant yet easily observable morphological traits, both derived from the neural crest, offered us an unparalleled opportunity to explore the differences in embryonic processes in the context of adult phenotypic variation.

Overall, the early development of the four Malawi cichlids presented here is characterised by many clade-generic features common to other teleosts (e.g. neurulation via cavitation of the neural keel) and, more specifically, other mouthbrooding cichlids, including the large maternal yolk supply and a direct development without a pronounced larval stage (Jones, 1972; Woltering et al., 2018). On the other hand, a careful comparison conducted in standardised conditions revealed a considerable degree of variability between these closely related species that contributes to their adult morphological diversity. This largely underexplored biological variation was primarily evident as anatomical and temporal (hetechronic) differences and was detected throughout early ontogeny, including the period of embryogenesis (e.g. somitogenesis) and post-hatching development (e.g. cartilage deposition and pigmentation development).

### 4.1 Heterochronies are common in cichlid development

One of the most prevalent inter- and intraspecific differences concerned the variability in the timing, rate and duration of specific developmental events. These included processes occurring in early embryonic development (i.e. segmentation) as well as later, throughout post-embryonic stages (e.g. formation of the chondrocranium). Although heterochronies in early development have been reported among cichlids (Kratochwil et al., 2015), including both within and between clutches of the same species (Morrison et al., 2001), at least some of this variation could have been attributed to the effects of temperature on the speed of development in teleost fishes (Schröter et al., 2008). Our results show that these differences persist in controlled conditions. Consequently, heterochronies might render the use of chronological age in cross-species comparative work potentially confounding, thus we recommend the use of both the morphological staging and the time elapsed from fertilisation in combination to infer developmental age and compare trait development and evolution.

The earliest and most striking differences were identified during somitogenesis, specifically in the number of generated somites in AC and RC, despite comparable total duration of the segmentation period. Intriguingly, the embryos did not visibly differ from one another in their progression of anatomical development when stage-matched by absolute somite number. Taken together, our results suggest that the modification of the somite ‘bauplan’, which can be considered as about 30-32 somites, was two-fold. It involved 1) acceleration of the somite clock throughout segmentation and 2) the formation of the additional few somites at the very end of this period (i.e. an increase in the total number of cycles). Following these observations, we hypothesise that, due to its key roles in the development and patterning of the animal body, variability in this fundamental morphogenetic process of embryogenesis could have an important yet largely underexplored function in the evolution of cichlid phenotypic diversity. Firstly, somites make broad contributions to the adult form, including the vertebral column and rib cage, cartilage and tendons, skeletal muscle and skin (Devoto et al., 2006; Holley, 2007; Morin-Kensicki et al., 2002; Stickney et al., 2000). The variation in the number of vertebrae has been previously described in the *Rhamphochromis* genus to range from 36 to 40 (Eccles and Trewavas, 1989), whereas *Astatotilapia burtoni* has 27 or 28 vertebrae (Woltering et al., 2018). Considering the relationship between the evolution of the skeletal system, including the vertebral column, and the body shape in the evolutionary history of vertebrates (Jones et al., 2018; Lindell, 1994; Müller et al., 2010; Richardson et al., 1998), it thus likely that divergence in the morphology of these derivatives contributes to the diversity of the body shapes across cichlid taxa (Malinsky et al., 2015). Secondly, the developmental dynamics of somitogenesis have been shown to influence the temporally coincident NC development, particularly in aspects such as the timing of delamination of NCCs from the neural tube and their subsequent migratory pathways (Loring and Erickson, 1987; Rocha et al., 2020; Sela-Donenfeld and Kalcheim, 2000; Teillet et al., 1987). Altogether, it is becoming clear that a good understanding of the processes of embryogenesis from comparative studies with wide taxon sampling will be required to elucidate its contribution to the developmental origins of morphological diversity of the clade.

### 4.2 The species-specific phenotypes of NC-derived traits are determined early in ontogeny

The stunning diversity of cichlid body pigmentation patterns and craniofacial shapes renders them a perfect model to study the genetic and developmental basis of phenotypic trait diversification. Our results show that both NC-derived features have visibly species-specific morphologies from the earliest stages of overt development.

It is possible that observed temporal differences in chromatophore appearance are related to the different contributions of each cell type to adult phenotype, for instance the earlier appearance of xanthophores in TM than in AC ‘Mbaka’ and RC could be linked to the differences in the extents of underlying yellow pigmentation in the adult colouration. The variation in timing could be also related to cell-cell interactions between individual cell types and their environment (Patterson and Parichy, 2019), with temporal shifts in the appearance of melanophores potentially influencing the subsequent emergence of xanthophores and iridophores. Further experimental work, including quantification of the chromatophore abundance, will be required to address these hypotheses.

Finally, the formation of body colouration in the examined head region did not seem to involve migration of mature, pigment-bearing cells but rather a sequential appearance of new chromatophores. This finding is in line with the account of colour pattern formation in Lake Malawi cichlids *Dimidiochromis compressiceps* and *Copadichromis azureus* (Hendrick et al., 2019) but unlike zebrafish, where pigmented melanophores are known to exhibit migratory behaviour, at least during stripe formation (Eom et al., 2012; Takahashi and Kondo, 2008). The species-specific trajectories of cichlid colouration could be thus set up prior to that, for instance during migration of the undifferentiated pigment cells through epidermis as suggested by Hendrick et al. (2019). Examination of the differentiation programme and migratory behaviour of the different chromatophore lineages could provide crucial insights into these questions.

Similarly to body pigmentation, the developmental and genetic basis of the vast variation in cichlid facial morphology has been previously investigated in several taxa (Albertson and Kocher, 2006; Conith et al., 2018; Kocher et al., 1993; Powder et al., 2014, 2015; Woltering et al., 2018). Our results expand this sampling by addition of three species occupying distinct positions on the ecomorphological axis, including the generalist, ancestor-like AC. Considering the vast temporal variability between our study species and in contrast to previous studies, we primarily focused on comparison of the developmental (stage), rather than chronological (dpf), stages of cartilage formation. The (overall) craniofacial ontogenies were conserved among sampled species with homologous elements forming in the same order.

The ontogeny was the primary axis of shape variation between species in both examined aspects of craniofacial shapes, indicating that the differences between developmental stages exceeded those present between species. Nonetheless, the morphological differences between species were detected from the onset of chondrogenesis in stage-wise comparisons. Notably, these revealed that both lateral and ventral aspects followed an interesting trend where initially distinct phenotypes (st. 18) converge towards one another (st. 19) to diverge again (st. 20) in morphospace.

Overall, our results are largely consistent with Powder et al. (2015) who also reported the ontogeny to be the main determinant of the cichlid craniofacial development in six Malawi species. Within that ontogenetic framework, based on comparisons using chronological age (expressed in dpf), the authors identified heterochronies as one of the drivers of species-specific phenotypes. By using a complementary approach of stage-wise comparisons, we provide evidence that the differences in craniofacial development exist as soon as cartilage forms, irrespective of the influence of heterochrony.

In conclusion, our results add further evidence that the divergence of developmental trajectories for both hyper-diverse cichlid morphologies of the craniofacial complex and skin pigmentation is specified during early embryonic development, prior to the overt formation of the trait. Moreover, the variability in developmental processes, exemplified here as heterochronic shifts in chromatophore appearance and morphogenesis of the chondrocranium, acts within conserved frameworks (e.g. following a conserved order of events) to contribute to species-specific phenotypes. The common embryonic origin of pigmentation and craniofacial phenotypes suggests that the differences observed at the onset of overt trait appearance may result from variation occurring during formation, migration and differentiation of the NC. In that scenario, we would expect that the sub-populations of the NC differentiating into the cartilage and pigment cell lineages to follow species-specific developmental trajectories, manifested for instance as differential migratory patterns of these cells, sizes of progenitor pools or heterochronies. Thus far, Powder et al. (2014) have reported that the alternate short and long jaw morphologies of *Labeotropheus fuelleborni* and *Maylandia zebra* (Lake Malawi), respectively, are associated with a non-synonymous mutation in the *limb bud and heart homolog* (*lbh*) gene. This mutation was demonstrated experimentally to result in altered migration patterns of the NCCs in *D. rerio* and *Xenopus*. Similarly, differential expression levels of *pax3a*, mediating xanthophore specification from the NC (Minchin and Hughes, 2008), have been implicated in continuous variation in colour patterns in *L. fuelleborni* and *Tropheops* ‘red cheek’ (Lake Malawi) (Albertson et al., 2014). Despite the compelling evidence suggesting the role of NC in the morphological diversity of cichlids, the innate patterns of NCC migration, as well as more general features of the NC development, remain to be explored in the cichlid system (Brandon et al., 2022).

## 5 Conclusions: the use of cichlids as a model for experimental evo-devo

Recent years have seen an increased interest in adopting cichlid fishes as an experimental model system. Despite the considerable advances, several aspects of fundamental cichlid biology, especially concerning embryonic development, have been largely overlooked. This study highlights the potential of East African cichlids as a valuable addition to the existing repertoire of teleost models, especially for research questions concerning evolution of embryogenesis, such as axial development and patterning, as well as variability in development of NC-derived traits. This variation, in turn, suggests that the differences in NC development may underlie the trait diversity. Considering the vast inter- and intraspecific diversity of cichlid fishes, we are certain that cichlid comparative embryology will provide important insights into vertebrate development and evolution.

## Acknowledgements

The authors thank Dr J. Andrew Gillis for help with histology and Dr Lewis Thomson for technical assistance with sample collection as well as the staff working in the animal facility in Madingley for maintenance of fish stocks. We thank Dr G. Turner (Bangor University) and Dr D. Joyce and Alan Smith (University of Hull) for the kind gifts of fish stocks. We also thank Bethan Clark for valuable comments on the manuscript and the Zoology Imaging Facility for assistance and support with microscopy. AM was supported by the Wellcome Trust PhD Programme in Developmental Mechanisms (215223/Z/19/Z). MES was supported by a Natural Environment Research Council Independent Research Fellowship (NE/R01504X/1).

## Contributions

AM and MES conceived the project and designed the experiments. AM, CY and SM performed the experiments and AM analysed the data. AM wrote the manuscript with contributions or feedback from all authors. All authors read and approved the final version of the manuscript.

## References

Alberch, P., Gould, S.J., Oster, G.F., and Wake, D.B. (1979). Size and Shape in Ontogeny and Phylogeny. Paleobiology 5, 296–317.

Albertson, R.C., and Kocher, T.D. (2006). Genetic and developmental basis of cichlid trophic diversity. Heredity 97, 211–221. https://doi.org/10.1038/sj.hdy.6800864.

Albertson, R.C., Powder, K.E., Hu, Y., Coyle, K.P., Roberts, R.B., and Parsons, K.J. (2014). Genetic basis of continuous variation in the levels and modular inheritance of pigmentation in cichlid fishes. Mol Ecol 23, 5135–5150. https://doi.org/10.1111/mec.12900.

Balon, E.K. (1977). Early ontogeny of Labeotropheus Ahl, 1927 (Mbuna, Cichlidae, Lake Malawi), with a discussion on advanced protective styles in fish reproduction and development. Environ Biol Fish 2, 147–176. https://doi.org/10.1007/BF00005370.

Brandon, A.A., Almeida, D., and Powder, K.E. (2022). Neural crest cells as a source of microevolutionary variation. Seminars in Cell & Developmental Biology https://doi.org/10.1016/j.semcdb.2022.06.001.

Bronner, M.E., and LeDouarin, N.M. (2012). Development and evolution of the neural crest: An overview. Developmental Biology 366, 2–9. https://doi.org/10.1016/j.ydbio.2011.12.042.

Bronner, M.E., and Simões-Costa, M. (2016). Chapter Seven - The Neural Crest Migrating into the Twenty-First Century. In Current Topics in Developmental Biology, P.M. Wassarman, ed. (Academic Press), pp. 115–134.

Brzozowski, F., Roscoe, J., Parsons, K., and Albertson, C. (2012). Sexually Dimorphic Levels of Color Trait Integration and the Resolution of Sexual Conflict in Lake Malawi Cichlids. Journal of Experimental Zoology Part B: Molecular and Developmental Evolution 318, 268–278. https://doi.org/10.1002/jez.b.22443.

Clark, B., Elkin, J., Marconi, A., Turner, G.F., Smith, A.M., Joyce, D., Miska, E.A., Juntti, S.A., and Santos, M.E. (2022). Oca2 targeting using CRISPR/Cas9 in the Malawi cichlid Astatotilapia calliptera. Royal Society Open Science 9, 220077. https://doi.org/10.1098/rsos.220077.

Conith, M.R., Hu, Y., Conith, A.J., Maginnis, M.A., Webb, J.F., and Albertson, R.C. (2018). Genetic and developmental origins of a unique foraging adaptation in a Lake Malawi cichlid genus. PNAS 115, 7063–7068. https://doi.org/10.1073/pnas.1719798115.

Devoto, S.H., Stoiber, W., Hammond, C.L., Steinbacher, P., Haslett, J.R., Barresi, M.J.F., Patterson, S.E., Adiarte, E.G., and Hughes, S.M. (2006). Generality of vertebrate developmental patterns: evidence for a dermomyotome in fish. Evolution & Development 8, 101–110. https://doi.org/10.1111/j.1525-142X.2006.05079.x.

Douarin, N.L., and Kalcheim, C. (1999). The Neural Crest (Cambridge University Press).

Eccles, D., and Trewavas, E. (1989). Malawian Cichlid Fishes the Classification of Some Haplochromine Genera (Germany: Herten).

Eom, D.S., Inoue, S., Patterson, L.B., Gordon, T.N., Slingwine, R., Kondo, S., Watanabe, M., and Parichy, D.M. (2012). Melanophore Migration and Survival during Zebrafish Adult Pigment Stripe Development Require the Immunoglobulin Superfamily Adhesion Molecule Igsf11. PLOS Genetics 8, e1002899. https://doi.org/10.1371/journal.pgen.1002899.

Fujimura, K., and Okada, N. (2007). Development of the embryo, larva and early juvenile of Nile tilapia Oreochromis niloticus (Pisces: Cichlidae). Developmental staging system. Development, Growth & Differentiation 49, 301–324. https://doi.org/10.1111/j.1440-169X.2007.00926.x.

Genner, M.J., and Turner, G.F. (2005). The mbuna cichlids of Lake Malawi: a model for rapid speciation and adaptive radiation. Fish and Fisheries 6, 1–34. https://doi.org/10.1111/j.1467-2679.2005.00173.x.

Hendrick, L.A., Carter, G.A., Hilbrands, E.H., Heubel, B.P., Schilling, T.F., and Le Pabic, P. (2019). Bar, stripe and spot development in sand-dwelling cichlids from Lake Malawi. EvoDevo 10, 18. https://doi.org/10.1186/s13227-019-0132-7.

Henning, F., and Meyer, A. (2014). The Evolutionary Genomics of Cichlid Fishes: Explosive Speciation and Adaptation in the Postgenomic Era. Annual Review of Genomics and Human Genetics 15, 417–441. https://doi.org/10.1146/annurev-genom-090413-025412.

Holley, S.A. (2007). The genetics and embryology of zebrafish metamerism. Developmental Dynamics 236, 1422–1449. https://doi.org/10.1002/dvdy.21162.

Hubaud, A., and Pourquié, O. (2014). Signalling dynamics in vertebrate segmentation. Nat Rev Mol Cell Biol 15, 709–721. https://doi.org/10.1038/nrm3891.

Jones, A.J. (1972). The early development of substrate-brooding cichlids (Teleostei: Cichlidae) with a discussion of a new system of staging. Journal of Morphology 136, 255–272. https://doi.org/10.1002/jmor.1051360302.

Jones, K.E., Angielczyk, K.D., Polly, P.D., Head, J.J., Fernandez, V., Lungmus, J.K., Tulga, S., and Pierce, S.E. (2018). Fossils reveal the complex evolutionary history of the mammalian regionalized spine. Science 361, 1249–1252. https://doi.org/10.1126/science.aar3126.

Jong, I.M.L. de, Witte, F., and Richardson, M.K. (2009). Developmental stages until hatching of the Lake Victoria cichlid Haplochromis piceatus (Teleostei: Cichlidae). Journal of Morphology 270, 519–535. https://doi.org/10.1002/jmor.10716.

Juntti, S.A., Hu, C.K., and Fernald, R.D. (2013). Tol2-Mediated Generation of a Transgenic Haplochromine Cichlid, Astatotilapia burtoni. PLoS ONE 8, e77647. https://doi.org/10.1371/journal.pone.0077647.

Kimmel, C.B., Ballard, W.W., Kimmel, S.R., Ullmann, B., and Schilling, T.F. (1995). Stages of embryonic development of the zebrafish. Developmental Dynamics 203, 253–310. https://doi.org/10.1002/aja.1002030302.

Klingenberg, C.P. (2011). MorphoJ: an integrated software package for geometric morphometrics. Mol Ecol Resour 11, 353–357. https://doi.org/10.1111/j.1755-0998.2010.02924.x.

Kocher, T.D. (2004). Adaptive evolution and explosive speciation: the cichlid fish model. Nature Reviews Genetics 5, 288–298. https://doi.org/10.1038/nrg1316.

Kocher, T.D., Conroy, J.A., McKaye, K.R., and Stauffer, J.R. (1993). Similar Morphologies of Cichlid Fish in Lakes Tanganyika and Malawi Are Due to Convergence. Molecular Phylogenetics and Evolution 2, 158–165. https://doi.org/10.1006/mpev.1993.1016.

Kratochwil, C.F., Sefton, M.M., and Meyer, A. (2015). Embryonic and larval development in the Midas cichlid fish species flock (Amphilophus spp.): a new evo-devo model for the investigation of adaptive novelties and species differences. BMC Dev Biol 15. https://doi.org/10.1186/s12861-015-0061-1.

Kratochwil, C.F., Liang, Y., Gerwin, J., Woltering, J.M., Urban, S., Henning, F., Machado-Schiaffino, G., Hulsey, C.D., and Meyer, A. (2018). Agouti-related peptide 2 facilitates convergent evolution of stripe patterns across cichlid fish radiations. Science 362, 457–460. https://doi.org/10.1126/science.aao6809.

Kratochwil, C.F., Kautt, A.F., Nater, A., Härer, A., Liang, Y., Henning, F., and Meyer, A. (2022). An intronic transposon insertion associates with a trans-species color polymorphism in Midas cichlid fishes. Nat Commun 13, 296. https://doi.org/10.1038/s41467-021-27685-8.

Li, C.-Y., Steighner, J.R., Sweatt, G., Thiele, T.R., and Juntti, S.A. (2021). Manipulation of the Tyrosinase gene permits improved CRISPR/Cas editing and neural imaging in cichlid fish. Sci Rep 11, 15138. https://doi.org/10.1038/s41598-021-94577-8.

Liang, Y., Gerwin, J., Meyer, A., and Kratochwil, C.F. (2020). Developmental and Cellular Basis of Vertical Bar Color Patterns in the East African Cichlid Fish Haplochromis latifasciatus. Front. Cell Dev. Biol. 8. https://doi.org/10.3389/fcell.2020.00062.

Lindell, L.E. (1994). The Evolution of Vertebral Number and Body Size in Snakes. Functional Ecology 8, 708–719. https://doi.org/10.2307/2390230.

Loh, Y.-H.E., Katz, L.S., Mims, M.C., Kocher, T.D., Yi, S.V., and Streelman, J.T. (2008). Comparative analysis reveals signatures of differentiation amid genomic polymorphism in Lake Malawi cichlids. Genome Biology 9, R113. https://doi.org/10.1186/gb-2008-9-7-r113.

Loring, J.F., and Erickson, C.A. (1987). Neural crest cell migratory pathways in the trunk of the chick embryo. Developmental Biology 121, 220–236. https://doi.org/10.1016/0012-1606(87)90154-0.

Lowery, L.A., and Sive, H. (2004). Strategies of vertebrate neurulation and a re-evaluation of teleost neural tube formation. Mechanisms of Development 121, 1189–1197. https://doi.org/10.1016/j.mod.2004.04.022.

Malinsky, M., Challis, R.J., Tyers, A.M., Schiffels, S., Terai, Y., Ngatunga, B.P., Miska, E.A., Durbin, R., Genner, M.J., and Turner, G.F. (2015). Genomic islands of speciation separate cichlid ecomorphs in an East African crater lake. Science 350, 1493–1498. https://doi.org/10.1126/science.aac9927.

Malinsky, M., Svardal, H., Tyers, A.M., Miska, E.A., Genner, M.J., Turner, G.F., and Durbin, R. (2018). Whole genome sequences of Malawi cichlids reveal multiple radiations interconnected by gene flow. BioRxiv 143859. https://doi.org/10.1101/143859.

McKinney, M.L., and McNamara, K.J. (1992). Heterochrony. The Evolution of Ontogeny. Geological Magazine - GEOL MAG 129. https://doi.org/10.1017/S0016756800021919.

Meyer, A. (1993). Phylogenetic relationships and evolutionary processes in East African cichlid fishes. Trends in Ecology & Evolution 8, 279–284. https://doi.org/10.1016/0169-5347(93)90255-N.

Meyer, A., Kocher, T.D., Basasibwaki, P., and Wilson, A.C. (1990). Monophyletic origin of Lake Victoria cichlid fishes suggested by mitochondrial DNA sequences. Nature 347, 550–553. https://doi.org/10.1038/347550a0.

Minchin, J.E.N., and Hughes, S.M. (2008). Sequential actions of Pax3 and Pax7 drive xanthophore development in zebrafish neural crest. Developmental Biology 317, 508–522. https://doi.org/10.1016/j.ydbio.2008.02.058.

Moran and Kornfield (1993). Retention of an Ancestral Polymorphism in the Mbuna Species Flock (Teleostei: Cichlidae) of Lake Malawi. Molecular Biology and Evolution https://doi.org/10.1093/oxfordjournals.molbev.a040063.

Morin-Kensicki, E.M., Melancon, E., and Eisen, J.S. (2002). Segmental relationship between somites and vertebral column in zebrafish. Development 129, 3851–3860. https://doi.org/10.1242/dev.129.16.3851.

Morrison, C.M., Miyake, T., and Wright, J.R. (2001). Histological study of the development of the embryo and early larva of Oreochromis niloticus (Pisces: Cichlidae). Journal of Morphology 247, 172–195. https://doi.org/10.1002/1097-4687(200102)247:2<172::AID-JMOR1011>3.0.CO;2-H.

Müller, J., Scheyer, T.M., Head, J.J., Barrett, P.M., Werneburg, I., Ericson, P.G.P., Pol, D., and Sánchez-Villagra, M.R. (2010). Homeotic effects, somitogenesis and the evolution of vertebral numbers in recent and fossil amniotes. Proceedings of the National Academy of Sciences 107, 2118–2123. https://doi.org/10.1073/pnas.0912622107.

Patterson, L.B., and Parichy, D.M. (2019). Zebrafish Pigment Pattern Formation: Insights into the Development and Evolution of Adult Form. Annual Review of Genetics 53. https://doi.org/10.1146/annurev-genet-112618-043741.

Powder, K.E., and Albertson, R.C. (2016). Cichlid fishes as a model to understand normal and clinical craniofacial variation. Developmental Biology 415, 338–346. https://doi.org/10.1016/j.ydbio.2015.12.018.

Powder, K.E., Cousin, H., McLinden, G.P., and Craig Albertson, R. (2014). A Nonsynonymous Mutation in the Transcriptional Regulator lbh Is Associated with Cichlid Craniofacial Adaptation and Neural Crest Cell Development. Mol Biol Evol 31, 3113–3124. https://doi.org/10.1093/molbev/msu267.

Powder, K.E., Milch, K., Asselin, G., and Albertson, R.C. (2015). Constraint and diversification of developmental trajectories in cichlid facial morphologies. EvoDevo 6, 25. https://doi.org/10.1186/s13227-015-0020-8.

Richardson, M.K., Allen, S.P., Wright, G.M., Raynaud, A., and Hanken, J. (1998). Somite number and vertebrate evolution. Development 125, 151–160. https://doi.org/10.1242/dev.125.2.151.

Roberts, R.B., Moore, E.C., and Kocher, T.D. (2017). An allelic series at pax7a is associated with colour polymorphism diversity in Lake Malawi cichlid fish. Molecular Ecology 26, 2625–2639. https://doi.org/10.1111/mec.13975.

Rocha, M., Singh, N., Ahsan, K., Beiriger, A., and Prince, V.E. (2020). Neural crest development: insights from the zebrafish. Developmental Dynamics 249, 88–111. https://doi.org/10.1002/dvdy.122.

Rohlf, F. (2010). TpsDig2: digitize coordinates of landmarks and capture outlines. Department of Ecology & Evolution, Stony Brook University, Stony Brook, NY, USA.

Saemi-Komsari, M., Salehi, M., Mansouri-Chorehi, M., Eagderi, S., and Mousavi-Sabet, H. (2018). Developmental morphology and growth patterns of laboratory-reared giraffe cichlid, Nimbochromis venustus Boulenger, 1908. International Journal of Aquatic Biology 6, 170–178. https://doi.org/10.22034/ijab.v6i3.525.

Salzburger, W. (2018). Understanding explosive diversification through cichlid fish genomics. Nat Rev Genet 19, 705–717. https://doi.org/10.1038/s41576-018-0043-9.

Santos, M.E., Braasch, I., Boileau, N., Meyer, B.S., Sauteur, L., Böhne, A., Belting, H.-G., Affolter, M., and Salzburger, W. (2014). The evolution of cichlid fish egg-spots is linked with a cis-regulatory change. Nat Commun 5, 5149. https://doi.org/10.1038/ncomms6149.

Schindelin, J., Arganda-Carreras, I., Frise, E., Kaynig, V., Longair, M., Pietzsch, T., Preibisch, S., Rueden, C., Saalfeld, S., Schmid, B., et al. (2012). Fiji: an open-source platform for biological-image analysis. Nat Methods 9, 676–682. https://doi.org/10.1038/nmeth.2019.

Schröter, C., Herrgen, L., Cardona, A., Brouhard, G.J., Feldman, B., and Oates, A.C. (2008). Dynamics of zebrafish somitogenesis. Developmental Dynamics 237, 545–553. https://doi.org/10.1002/dvdy.21458.

Sela-Donenfeld, D., and Kalcheim, C. (2000). Inhibition of noggin expression in the dorsal neural tube by somitogenesis: a mechanism for coordinating the timing of neural crest emigration. Development 127, 4845–4854. https://doi.org/10.1242/dev.127.22.4845.

Stickney, H.L., Barresi, M.J.F., and Devoto, S.H. (2000). Somite development in zebrafish. Developmental Dynamics 219, 287–303. https://doi.org/10.1002/1097-0177(2000)9999:9999<::AID-DVDY1065>3.0.CO;2-A.

Takahashi, G., and Kondo, S. (2008). Melanophores in the stripes of adult zebrafish do not have the nature to gather, but disperse when they have the space to move. Pigment Cell & Melanoma Research 21, 677–686. https://doi.org/10.1111/j.1755-148X.2008.00504.x.

Teillet, M.-A., Kalcheim, C., and Le Douarin, N.M. (1987). Formation of the dorsal root ganglia in the avian embryo: Segmental origin and migratory behavior of neural crest progenitor cells. Developmental Biology 120, 329–347. https://doi.org/10.1016/0012-1606(87)90236-3.

Woltering, J.M., Holzem, M., Schneider, R.F., Nanos, V., and Meyer, A. (2018). The skeletal ontogeny of Astatotilapia burtoni – a direct-developing model system for the evolution and development of the teleost body plan. BMC Developmental Biology 18, 8. https://doi.org/10.1186/s12861-018-0166-4.S

