## Supporting information for "Morphological and temporal variation in early embryogenesis contributes to species divergence in Malawi cichlid fishes"

**Supplementary Table S1. Summary of the stages and corresponding developmental landmarks referenced throughout this study, adapted from the staging table of Nile tilapia *Oreochromis niloticus*** **(Fujimura and Okada, 2007)** **with modifications to reflect features unique to examined species.** It covers the period from zygote up to a free-feeding early juvenile (i.e. until the abdominal body wall closure is complete, about 3 weeks post fertilisation) and includes 27 or 28 stages corresponding to defined developmental characteristics that are usually easily identifiable under a simple compound microscope such as development of the eye pigmentation (for late embryonic stages) or the number of elements in the central caudal fin ray (applicable for the stages post-hatching and thereafter abbreviated as CFRE (Otten, 1981)). Timings for Malawi cichlids as given for *Astatotilapia calliptera*.

| **Nile tilapia (*Orechromis niloticus*)** **(Fujimura and Okada, 2007)** | | | | | | | **This study** | | |
| --- | --- | --- | --- | --- | --- | --- | --- | --- | --- |
| Period | | Stage | hpf^†^ | dpf^†^ | Characteristics | Stage description | Developmental landmarks used to define the stage | hpf in AC | dpf in AC^‡^ |
| Embryo | Zygote | 1 | 0-1.5 |  | 1-cell |  |  | 0 |  |
|  | Cleavage | 2 | 1.5-2 |  | 2-cell |  |  | 2 |  |
|  |  | 3 | 2 |  | 4-cell |  |  | 4 |  |
|  |  | 4 | 3 |  | 8-cell |  |  | 6 |  |
|  |  | 5 | 4 |  | 16-cell |  |  | 8 |  |
|  | Blastula | 6 | 4-12 |  | Early blastula |  | Blastodisc (located on the animal pole of the yolk) has a spherical shape | 16 |  |
|  |  | 7 | 12-17 |  | Middle blastula |  | Blastodisc becomes elliptical | 20 |  |
|  |  | 8 | 17-22 |  | Late blastula |  | Blastodisc flattens | 26 | 1 |
|  | Gastrula | 9 | 22-26 | 2 | Gastrula, epiboly 30-50% |  | Epiboly begins – blastoderm resembles an inverted cup of uniform thickness | 30 | 1 |
|  | Segmentation | 10 | 26-30 | 2 | Neurula, epiboly 50-90%^§^ | The embryo thickens, especially at the anterior end. The brain primordium is visible. | Brain rudiment, <10 somites | 34-38 | 1 |
|  |  | 11 | 30-40 | 2-3 | Yolk plug closure and segmentation | The formation of the optic primordia; appearance of melanophores on yolk; prominent tailbud | Optic primordia, 10-17 somites | 38-48 | 1-2 |
|  |  | 12 | 40-44 | 3 | Optic cups and tail undercut | Melanophores on yolk become more prominent. The primordium of the brain begins to undergo regionalisation, forming the forebrain, midbrain, and hindbrain. The optic primordia develop optic cups and the otic placodes hollow, forming the otic vesicles. | Optic cups with lens, otic vesicles, brain ventricle, 18-24 somites | 48-56 | 2 |
|  |  | 13 | 44-48 | 3 | Brain differentiation | The brain thickened and visibly partitioned into the forebrain, midbrain, and hindbrain; The ventricle of the brain was also visible. The olfactory placodes were prominent on the surface anterior to the optic primordia with the lens placodes. The otic vesicle contained two otoliths that were tiny at this time. | Tripartite brain, olfactory placode, >24 somites | 54-72 | 2-3 |
|  | Pharyngula | 14 | 48-60 | 3-4 | Heartbeat | The embryo has approximately 32 somites. The optic cups with lens placodes and olfactory placodes are discernible. The median fin fold and anus begin to form. | Onset of eye pigmentation |  | 3 (late) |
|  |  | 15 | 60-72 | 4 | Onset of blood circulation | Ovoid otic vesicles form, each with two otoliths. The blood circulation starts as the heart differentiates with expansion of the pericardial cavity. Melanophores are present on the ventral side of the embryo along the anterior vitelline vein. | Increase in eye pigmentation, visible heartbeat |  | 4 (early) |
|  |  | 16 |  | 4-5 | Head enlargement | The optic tectum is differentiated in the midbrain. | Eyes are fully opaque, head has enlarged and rounded posteriorly |  | 4 (late) |
|  | Hatching | 17 |  | 5-6 | Jaw extension | Mouth begins to open, and embryos usually hatch. | Hatching followed by straightening of the body axis, mouth opening visible |  | 5 (early) |
|  |  | 18 |  | 6 | Gill formation | The midbrain protrudes dorsally as the position at the epiphyseal bar becomes depressed (visible from the lateral view). The blood circulation expands in the caudal fin and the fin rays begin to form in the hypural region. Numerous melanophores are present on the dorsal surface of the head. | Mouth opens and moves sporadically; the lower jaw is protruding; blood circulation visible in the caudal fin |  | 5 (late) |
| Larva^¶^ | Early larva | 19 |  | 6-7 | Opercular movement | Movements of the gills and the mouth are much stronger, and the mesenchymal condensations of dorsal and anal fins appear. | The lower jaws and gill covers are moving rapidly, mesenchymal condensations of dorsal and anal fins |  | 5-6 |
|  |  | 20 |  | 7 | 2 CFRE |  |  |  | 6 |
|  |  | 21 |  | 7-8 | 3 CFRE |  |  |  | 8 |
|  |  | 22 |  | 8-9 | 4 CFRE |  |  |  | 9 |
|  | Late larva | 23 |  | 9-10 | 5 CFRE |  |  |  | 10 |
|  |  | 24 |  | 9-10 | 6 CFRE |  |  |  | 12-13 |
|  |  | 25 |  | 11-13 | 7 CFRE |  |  |  | 13-14 |
| Juvenile |  | 26 |  | 12-13 | 8 CFRE |  |  |  | 15-16 |
|  |  | 27 |  | 14-15 | 9 CFRE |  |  |  | 16-18 |
|  |  | 28 |  |  | 10 CFRE |  | **Onset of juvenile period**^††^ | | |
|  |  | 29 |  |  | 11 CFRE |  |  | | |
|  |  | 30 |  |  | 12 CFRE |  |  |  |  |
|  |  | 31 |  |  | 13 CFRE |  |  |  |  |
|  |  | 32 |  |  | 14 CFRE |  |  |  |  |

AC – *Astatotilapia calliptera*; CFRE – caudal fin ray elements; dpf – days post-fertilisation; hpf – hours post-fertilisation.

†Rearing temperature 27 ^o^C.

‡Note that the day of fertilisation is denoted as 1 dpf, whereas in our study, it is 0 dpf.

§At this stage, embryos examined in our study remained at 30% epiboly.

¶In contrast to Fujimura and Okada, we do not distinguish period of larval development in examined cichlid embryos and as reported first in *Astatotilapia burtoni* (Woltering)

††Except for TM for which st. 28 (10 CFRE) is the last stage in the guide.

**Supplementary Table S2.** Sample number for species used in examination of developmental trajectories during somitogenesis and post-hatching stages (Figure X). For somitogenesis, n represents the total number of examined embryos, each at a single time point, whereas for post-hatching stages, n corresponds to the number of embryos followed and inspected daily from st. 16 until free-feeding stage. Clu - number of clutches from which the examined animals were sampled from.

| **Period** | **Species** | **Colour in figure** | **Sample size** |
| --- | --- | --- | --- |
| Somitogenesis (st. 10-13, Figure 8a-b) | *Astatotilapia calliptera* ‘Mbaka’ | 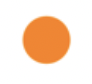 | N = 41  Clu = 5 |
|  | *Rhamphochromis* sp. ‘chilingali’ | 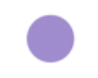 | N = 44  Clu = 3 |
| Post-hatching (st. 16-28, Figure 8c) | *Astatotilapia calliptera* ‘Mbaka’ | 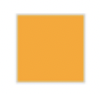 | N = 8  Clu = 3 |
|  | *Astatotilapia calliptera* ‘Salima’ | 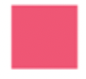 | N = 6  Clu = 3 |
|  | *Rhamphochromis* sp. ‘chilingali’ | 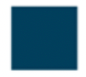 | N = 9  Clu = 3 |
|  | *Tropheops* sp. ‘mauve’ | 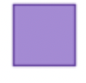 | N = 6  Clu = 3 |

**Supplementary Table S3. Sample number for species used in morphometric analyses.**

| **Species** | **Stage 18** | **Stage 19** | **Stage 20** |
| --- | --- | --- | --- |
| *Astatotilapia calliptera* | 8 | 15 | 11 |
| *Tropheops* sp. ‘mauve’ | 6 | 8 | 16 |
| *Rhamphochromis* sp. ‘chilingali’ | 12 | 14 | 10 |

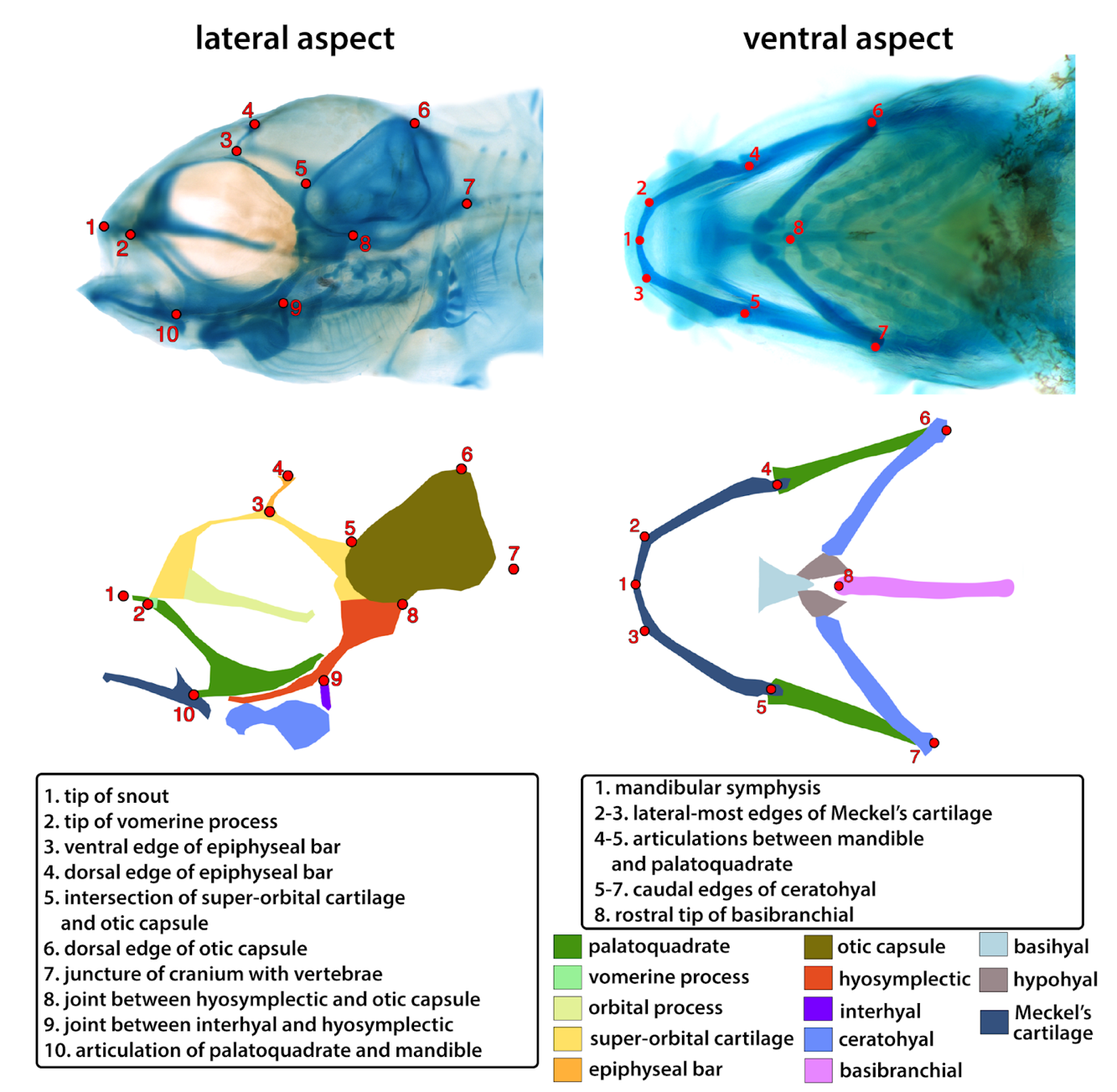

**Supplementary Figure S1. The landmarking protocol adapted from** **(Powder et al., 2015)****.** The landmarks 8, 9 and 11 from the original protocol were excluded from ventral analysis to account for the absence of specific anatomical structures at stage 18. For the lateral aspect, no semilandmarks between landmarks 2 and 3 were included.

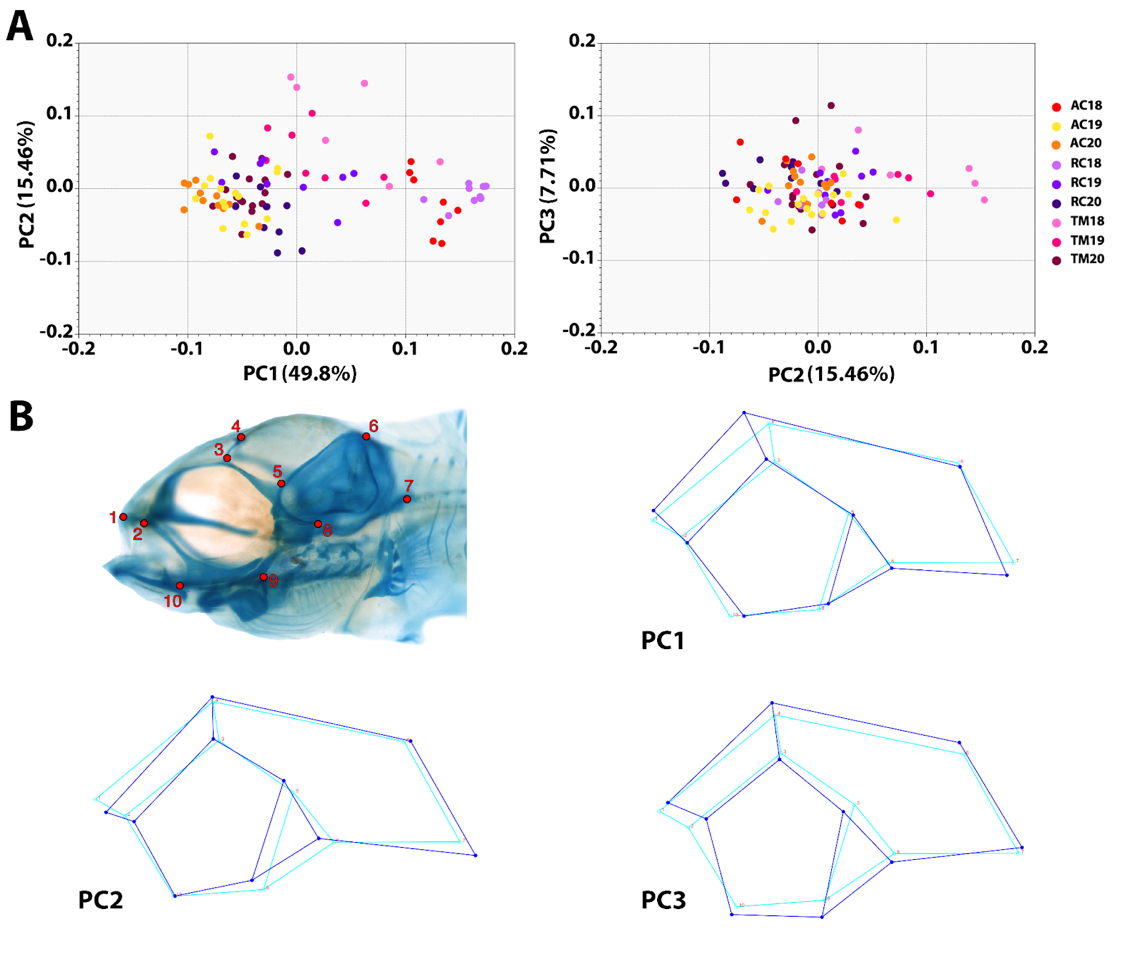

**Supplementary Figure S2. Principal Component Analysis (A) and wireframes (B) for lateral development of the craniofacial skeleton.** (A) The first three principal components (PCs) accounted for most of the shape variability (72.97%). The primary axis of variation (PC1, 49.8%) described differences in the craniofacial slope inferred from the changes to the slope of the line connecting the dorsal most point of the epiphyseal bar (LM4) to the rostral edge of the snout (LM1). PC2 (15.46%) accounted for the anterior outgrowth of the neurocranium (LMs 1 and 2) and the relative proportions between different elements of the chondrocranium (LM5, 7-10). Finally, PC3 (7.71%) described the outgrowth in the posterior (LM6) and facial regions (LMs 2 and 9). Colour of the points in A corresponds to the species and stages (18-20) as indicated in the legend on the right hand side. (B) Position of landmarks as used for geometric morphometric analysis of the lateral aspect. AC - *Astatotilapia calliptera*; PC - principal component; TM - *Tropheops* sp. ‘mauve’; RC - *Rhamphochromis* sp. ‘chilingali’.

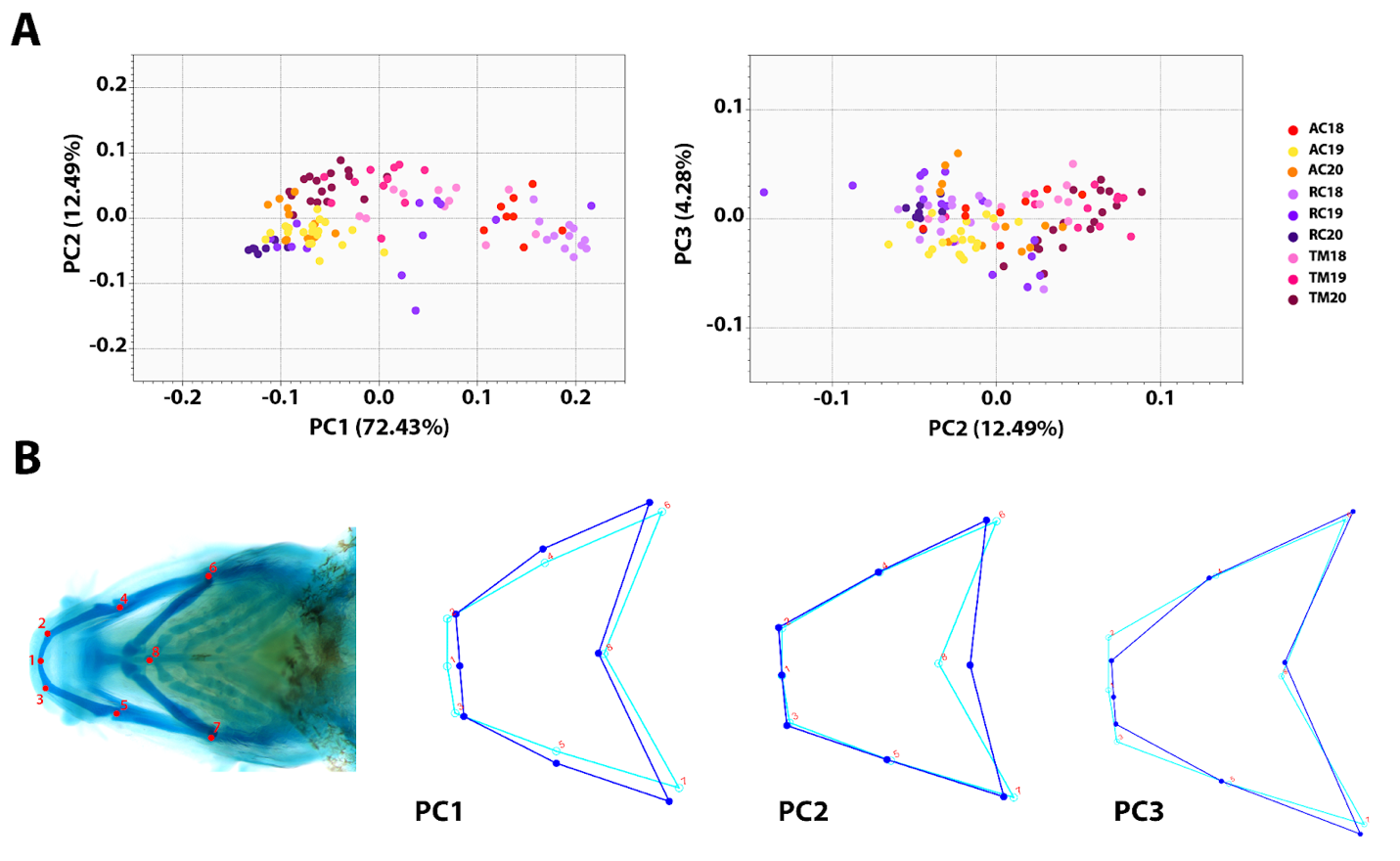

**Supplementary Figure S3. Principal Component Analysis (A) and wireframes (B) for ventral development of the craniofacial skeleton. (A)** Similarly to lateral development, in a dimensionality reduction analysis, the majority of shape variation (89.19%) was explained by PCs 1-3. PC1 (72.43%) described the width of the ventral aspect, particularly of the Meckel’s cartilage and pharyngeal skeleton (distances between LMs 2-3, LMs 4-5 and LMs 6-7) as well as the mandible length (distance from LM2 to LM4 and from LM3 to LM5). The second axis (PC2) accounted for 12.49% of total shape variation and corresponded to the position of the rostral tip of the basibranchial (LM8). PC3 (4.28%) described the width of the anterior Meckel’s cartilage (distances between LM2 and LM3 as well as between LMs 2-4 and LMs 3-5), reflecting the narrowing of the jaws in the anterior region. Colour of the points in A corresponds to the species and stages (18-20) as indicated in the legend on the right hand side. (B) Position of landmarks as used for geometric morphometric analysis of the ventral aspect. AC - *Astatotilapia calliptera*; PC - principal component; TM - *Tropheops* sp. ‘mauve’; RC - *Rhamphochromis* sp. ‘chilingali’.
